# Cannabinoid receptor 1 is required for neurodevelopment of striosome-dendron bouquets

**DOI:** 10.1101/2022.01.29.478320

**Authors:** Jill R. Crittenden, Tomoko Yoshida, Samitha Venu, Ara Mahar, Ann M. Graybiel

## Abstract

Cannabinoid receptor 1 (CB1R) has strong effects on neurogenesis and axon pathfinding in the prenatal brain. Endocannabinoids that activate CB1R are abundant in the early postnatal brain and in mother’s milk, but few studies have investigated their function in newborns. We examined postnatal CB1R expression in the major striatonigral circuit from striosomes of the striatum to the dopamine-containing neurons of the substantia nigra. CB1R enrichment was first detectable between postnatal days 5 and 7, and this timing coincided with the formation of ‘striosome-dendron bouquets’, the elaborate anatomical structures by which striosomal neurons control dopaminergic cell activity through inhibitory synapses. In *Cnr1^−/−^* knockouts lacking CB1R expression, striosome-dendron bouquets were markedly disorganized by postnatal day 11 and at adulthood, suggesting a postnatal pathfinding connectivity function for CB1R in connecting striosomal axons and dopaminergic neurons analogous to CB1R’s prenatal function in other brain regions. Our finding that CB1R plays a major role in postnatal wiring of the striatonigral dopamine system, with lasting consequences at least in mice, points to a crucial need to determine whether lactating mothers’ use of CB1R agonists (e.g., in marijuana) or antagonists (e.g., type 2 diabetes therapies) can disrupt brain development in nursing offspring.

## Introduction

Drugs of abuse augment levels of dopamine in striatal forebrain regions and motivate action sequences that can become extreme habits (Nelson & Killcross, 2006; Graybiel, 2008; Everitt, 2014; Gremel & Lovinger, 2017). Among the targets of these drugs are dopamine-containing neurons in the substantia nigra pars compacta (SNc). These neurons give rise to the nigrostriatal tract that degenerates in Parkinson’s disease, and they are vulnerable to abnormality in a range of motor and neuropsychiatric disorders. The nigrostriatal tract is known in the normal state to modulate not only movement but also psychic vigor, motivation and reinforcement-based learning.

Within the striatum, specialized widely distributed zones known as striosomes (aka patches) respond to psychomotor stimulants such as amphetamine by increased expression of immediate-early response genes (Graybiel et.al., 1990; Moratalla et al., 1996; Jedynak et al., 2012; Canales & Graybiel, 2000; Crittenden et al., 2017; Prager & Plotkin, 2019; Crittenden et al., 2021a). Axons from projection neurons in striosomes are enriched in cannabinoid receptor 1 (CB1R) and innervate clusters of dopamine-containing neurons in the ventral tier of the SNc (SNcv) to form anatomically conspicuous ‘striosome-dendron bouquets’ (Davis et al., 2018). Striosomal axons innervate the SNcv and also intertwine themselves in synapse-rich bundles of dopamine-containing dendrites (dendrons) that protrude deep into the substantia nigra pars reticulata (SNr), a major basal ganglia output nucleus (Crittenden et al., 2016). It now has been shown that optogenetic stimulation of striosomal axons within striosome-dendron bouquets can fully inhibit spike activity of dopamine-containing neurons (Lee & Tepper, 2009; McGregor et al., 2019; Evans et al., 2020) and further can produce rebound activation of the dopamine neurons (Evans et al., 2020).

Endocannabinoids as well as therapeutic and recreational cannabinoids bind to the receptors CB1R and CB2R (Ashton et al., 2008). CB1R is thought to mediate most of the psychoactive and habit-forming qualities of marijuana (Covey et al., 2017; Zimmer et al., 1999; but see Liu et al., 2017). Together, this evidence strongly suggests that CB1Rs, along with other receptors expressed in the bouquets such as the mu opioid receptor (MOR1; Crittenden et al., 2016), could have marked effects on neural circuit function, behavior and addiction by controlling the activity of dopamine-containing neurons (Davis et al., 2018; Riegel & Lupica, 2004). Functions of striosome-dendron bouquets in adults, however, are yet to be reported.

Here we demonstrate that striosome-dendron bouquets form in the early postnatal period, coinciding with the enrichment of CB1R in striosomal neurons and their axons. Moreover, mice lacking CB1R expression have abnormal placement of striosomal axons and dopaminergic dendrites such that the bouquets appsear disorganized already in the early postnatal period and in adulthood, indicating that CB1R signaling in the postnatal brain controls the wiring of the striosome-nigral circuit.

## Materials and Methods

### Mice

All animal procedures were approved by the animal care committee at Massachusetts Institute of Technology (MIT), which is AALAC accredited. Frozen embryonic heterozygous *Cnr1^−/+^* mice (B6.129-Cnr1<tm1Zim>/Ieg; #EM:02274, RRID: 3795243), created by Dr. A. Zimmer (Zimmer et al., 1999), were imported from the European Mouse Mutant Archive (EMMA) to MIT to establish a colony. Mice carrying the *P172-mCitrine* striosomal transgene marker (Piggyback-Tta, Line P172 C57B6J, http://enhancertrap.bio.brandeis.edu/data/) (Shima et al., 2016) were imported to MIT to establish a colony as previously described (Crittenden et al., 2016). Knockout (KO) and sibling control pups were generated from pairs of *Cnr1*^*−/+*^ heterozygous mice. In cases where the *P172-mCitrine* transgene was used, *Cnr1*^*−/+*^ heterozygous mice were crossed to *P172-mCitrine* hemizygous mice, and *Cnr1*^*−/+*^ offsprings carrying the *P172-mCitrine* transgene were then crossed to *Cnr1*^*−/+*^ mice to generate *Cnr1*^*-/-*^ KOs and *Cnr1*^*+/+*^ controls that each carry one copy of the *P172-mCitrine* transgene. Mice were genotyped by Transnetyx, Inc. for *neomycin* (present at the *Cnr1* deletion site), the wildtype *Cnr1* allele, and *TRE* (*tetracycline-responsive element* present at the *P172-mCitrine* transgene site).

Mice were group-housed, had free access to food and water and were maintained on a 12/12-h light/dark cycle (lights on at 7:00 AM). Roughly equal numbers of each sex were used for all experiments. Mice imaged for *P172-mCitrine* expression were evaluated at 5.5 weeks of age or younger because of age-dependent silencing of the *P172-mCitrine* transgene (Shima et al., 2016). Exceptions were 70 day old *P172-mCitrine* mice that were used for circuit tracing, as described in the Results. *Cnr1*^*-/-*^ knockout mice of both sexes were compared and no gross differences between sexes were evident for the described brain phenotype.

### Tissue Preparation and Immunoreactions

For a detailed protocol, see Crittenden et al. 2021b. For brain collection from adults, mice were deeply anesthetized with Euthasol (pentobarbital sodium and phenytoin sodium from Virbac AH Inc.) prior to transcardial perfusion with 20 ml of 0.9% saline and 60 ml of freshly depolymerized 4% paraformaldehyde in 0.1 M NaKPO4 buffer. Brains were then dissected, post-fixed for 90 min and stored in 25% glycerol sinking solution overnight or until cutting. For whole mount preparations, mice were perfused with heparin prior to fixative in order to eliminate fluorescence from red blood cells.

For brain tissue collection from pups, neonatal mice (P0-P5) were anesthetized on ice and older pups (P7, P11) by isoflurane inhalation exposure. After removing the scalp, the whole head was submerged in freshly depolymerized 4% paraformaldehyde in 0.1 M NaKPO4 buffer and placed on a low-speed shaker for 3 days at 4°C. The tissue was then transferred to a 25% glycerin solution and shaken overnight. Brain tissue was then dissected from the skull and returned to 25% glycerin until cutting.

To prepare sections, brains were frozen on dry ice to generate 30 μm (adult samples) or 60 μm (pup samples) free-floating coronal sections with a freezing microtome. Sections were stored in 0.1% sodium azide in 0.1 M NaKPO4 solution.

For the immunoreactions, sections were rinsed 3 times for 5 min each with 10 mM NaPO4, 150mM NaCl, 2.7 mM KCl with 0.2% Triton X-100 and then blocked by shaking for 60 min in TSA Blocking Reagent (Perkin Elmer). Sections were left shaking for 1-3 nights at 4°C in primary antibodies. Primary antibodies and concentrations used were: goat anti-CB1R (Frontier Institute CB1-Go-Af450, RRID: AB_2571592), 1:100; rat anti-DAT (Millipore MAB369, RRID: AB_2190413), 1:200; rabbit anti-TH (Abcam ab112, RRID: AB_297840), 1:4000; chicken anti-GFP (to detect mCitrine; Abcam ab13970, RRID: AB_300798), 1:2,000; rabbit anti-MOR1; Abcam Ab134054), 1:500; rabbit anti-FoxP2 (Sigma Aldrich HPA-000382, RRID: AB_1078908), 1:1,000. After rinsing, secondaries were applied at a 1:300 dilution for an overnight reaction at 4°C with gentle shaking. Secondary antibodies were: anti-goat Alexa fluor 647, (Thermo Fisher A21447, RRID: AB_2535864); anti-rat Alexa fluor 546 (Thermo Fisher A11081, RRID: AB_2534125), anti-rabbit Alexa fluor 546 (Thermo Fisher A10040, RRID: AB_2534016); anti-chicken FITC (Abcam ab63507, RRID: AB_1139472). Sections were mounted on subbed glass slides and coverslipped using ProLong Antifade Reagent with DAPI (Thermo Fisher Scientific).

### Whole Brain Clearing

Perfused brains were bisected and trimmed to contain mainly the basal ganglia. Brains were cleared using the uDISCO method (Pan et al., 2016) as follows. Samples were dehydrated by gentle shaking in increasing concentrations of *tert*-butanol in dH2O at room temperature: 30% for 4 h, 50% for 4 h, 70% overnight, 80% for 4 h, 90% for 4 h, and 96% overnight. Samples were then incubated in dichloromethane for 50 min and moved into a solution of BABB-D10 (1:2 ratio of benzyl alcohol and benzyl benzoate (BABB)), 10:1 ratio of BABB and diphenyl ether, 0.4% vol vitamin E for overnight incubation and further storage until imaging. All incubation steps were performed at room temperature under a fume hood. For confocal imaging, samples were transferred into BABB-D10 filled chambers with 1.5 coverslip-equivalent bottoms (ibidi Inc.).

### Microscopy

For fluorescence microscopy, a Zeiss AxioZoom microscope was used to obtain wide-field images with standard epifluorescence filter sets for DAPI (365 excitation, 395 beamsplitter, 445/50 emission), eGFP/AF488 (470/40 excitation, 495 beam splitter, 525/50 emission), tdTomato/AF546 (550/25 excitation, 570 beamsplitter, 605/70 emission) and Cy5/AF647 (640/30 excitation, 660 beamsplitter, 690/50 emission). Confocal imaging on sections and whole mount tissue was performed with a Zeiss LSM710 with diode lasers (473, 559, 653 nm) for excitation. Images were collected with a 10x 0.4 NA objective and a 60X oil 1.3 NA objective lens. Optical sectioning was optimized according to the microscope software. Images were processed and analyzed with Fiji software (Schindelin et al., 2012; RRID: SCR_002285) or, for 3-D reconstruction and movies, Imaris 9.5 software (Oxford Instruments; RRID: SCR_007370). Figures were prepared with Photoshop 22.4.2 and Illustrator 6.0 (Adobe).

### Histological Measurements

For striosomal counts, for striosomal and matrix compartment area calculations, and for average MOR1 immunointensity calculations in striosomes and matrix, measurements were made from images acquired on a Zeiss AxioZoom microscope. Each image was converted into 8-bit and analyzed with Fiji software. Fluorescence intensities were normalized by subtracting background fluorescence for each sample. A genotype blind investigator used the freehand selection tool to draw the boundaries of striosomes using MOR1 immunolabeling. Any striosomes that were connected were counted as one unit. Mean intensity value and pixel area were measured for each striosome and for the whole surrounding matrix.

Striosome-dendron bouquet and SNcv size measurements were made from images acquired on a Zeiss AxioZoom microscope and analyzed using Photoshop 23.1 (Adobe) and Fiji software. Each image was first converted into 8-bit, and fluorescence intensities were normalized by subtracting background fluorescence for each sample. Thresholding was used to create defined borders for dendrons. Measurements were made by investigators blinded to genotype. Using both the DAT and MOR1 channel, dendron widths (medial-lateral aspect) were measured using the ruler tool at two different distances, 50 μm and 100 μm, ventral to the SNcv/SNr border. SNcv regions were defined as double-labeled for DAT and MOR1, and depths (dorsal-ventral aspect) were measured at sites adjacent to dendrons.

### Cell Counting

Coronal brain sections at the anterior, mid-level and posterior striatum were labeled with DAPI and for immunodetection of P172-mCitrine and FoxP2 and imaged on a Zeiss Axiozoom microscope. Sections from *Cnr1*^*-/-*^ KO and sibling control mice were age-matched (27-33 days old range) and samples from each sex were taken and processed in the same way and in parallel by individuals blinded to genotype. In images of sections immunolabeled for MOR1, the striatum and striosomes were outlined by hand and P172-mCitrine-and FoxP2-positive cells within striosomes of the dorsal striatum (dorsal to the anterior commissure) were counted in each section. The entire area of the striatum was measured to calculate the overall density of P172-mCitrine-positive and FoxP2-positive striosomal neurons.

Identification of striosomes and immunopositive cells was assisted by Fiji software. The images were converted to 16 bit gray levels and inverted. The dorsal striatal area was selected, background signal was subtracted using the “Rolling Ball” command (radius = 500 μm), the “Analyze Particles” command was applied and verified by eye to be identifying bona fide cells, and the P172-mCitrine and FoxP2-positive cell counts were recorded.

### Statistics

Histological data measurements from multiple sections for each adult mouse (three striatal and six nigral hemisphere sections per mouse) were averaged to obtain one value per hemisphere per mouse. Each animal’s average value per hemisphere was plotted and used for standard deviation calculations in order to reflect inter-animal variability. Statistical differences between groups were assessed by unpaired, two-tailed Student’s t-tests. Specific *p* values are given when *p* < 0.05 and considered statistically significant.

## Results

### Striosome-dendron bouquets in the substantia nigra appeared severely disorganized in *Cnr1*^*-/-*^ KO mice

As a first test of CB1R function in striosomes and their axonal projections to bouquets, we examined brains of *Cnr1*^*-/-*^ KO mice (Zimmer et al., 1999) that were crossed to transgenic mice carrying a striosome-enriched flurophore marker, P172-mCitrine (Crittenden et al., 2016; Shima et al., 2016). *P172-mCitrine* transgenic mice were generated as part of a project in which the PiggyBac transposon was employed to insert the fluorophore-encoding *mCitrine* transgene at many different loci throughout the genome to yield mouse lines that label a variety of neuronal cell types. The *P172-mCitrine* line was discovered to drive highly specific expression in striosomal projection neurons that target striosome-dendron bouquets (Crittenden et al., 2016; Shima et al., 2016).

We imaged the entire basal ganglia in cleared whole brains from *Cnr1*^*-/-*^ KO and control mice carrying the *P172-mCitrine* transgene, using confocal fluorescence imaging and 3D reconstruction (**Fig. 1**). Similar to their sibling control mice, *Cnr1*^*-/-*^ KO mice had striosomes with axons that project toward their intended nigral target in the midbrain (**Fig. 1*A***). However, within the SN there were severe abnormalities in the distribution of striosomal fibers (**Fig. 1*B***). Instead of forming discrete striosome-dendron bouquet “stems” as in the controls, the P172-mCitrine striosomal fibers in *Cnr1*^*-/-*^ KO mice appeared in a thick pile-up at the presumed SNcv/SNr border where dopaminergic cell bodies of the SN ventral tier reside (Crittenden et al., 2016). The appearance of striosome-dendron bouquets in brain sections depends on the level and plane of section. To ensure that our impression of bouquet differences between *Cnr1*^*-/-*^ KOs and controls was not simply a reflection of this variability, we used confocal optical sectioning and 3D reconstruction from blocks of tissue that contained the entire basal ganglia to create 3D movies of striosomal neurons and their projections. These movies show that striosomal bouquet fibers were indeed grossly disorganized in *Cnr1*^*-/-*^ KO mice (**Movies 1, 2**).

**Figure 1.**
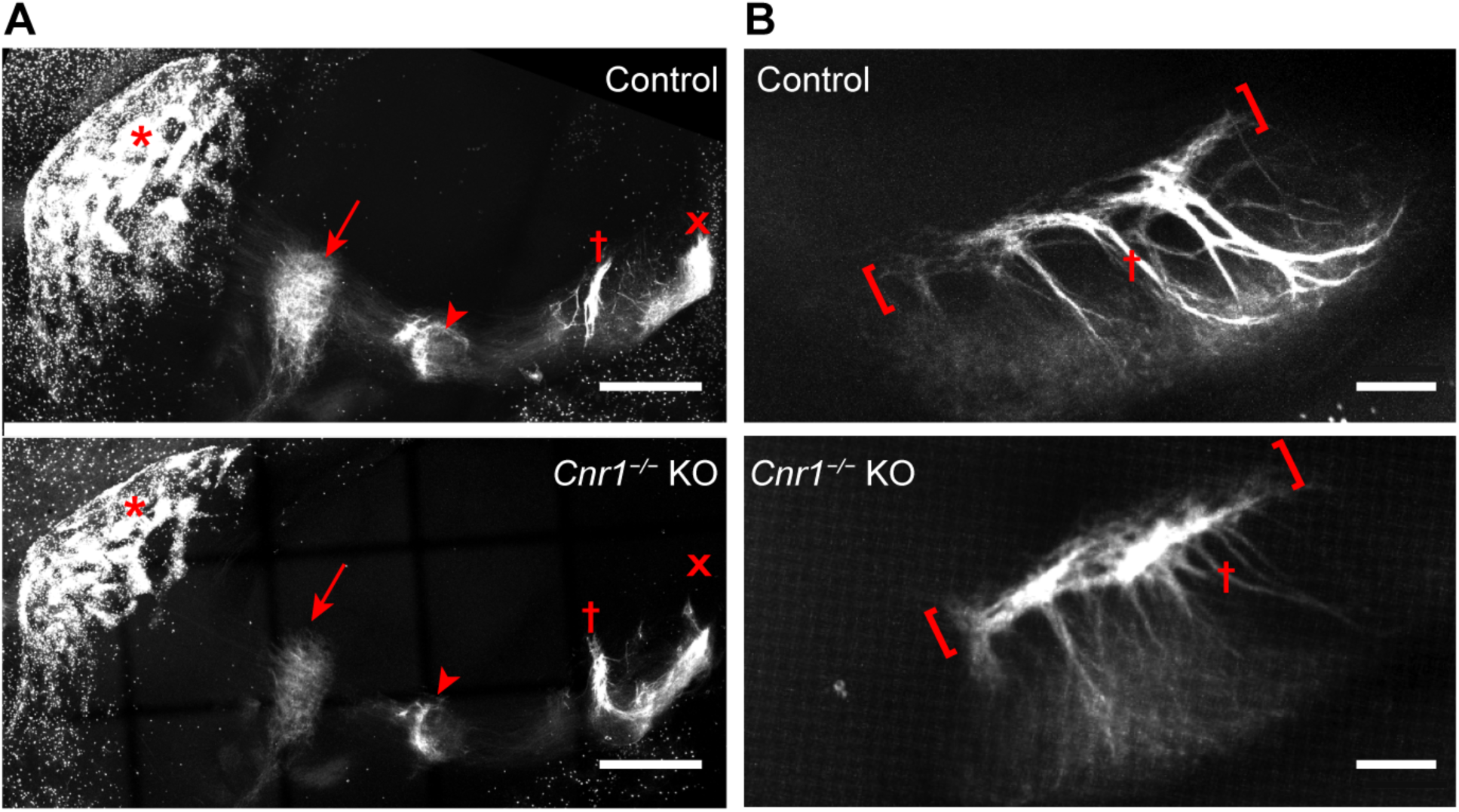
Striosomal axons reach their targets but striosome-dendron bouquets are severely malformed in the SN of *Cnr1*^*-/-*^ KO mice. To visualize the entire striosome-nigral projection, confocal imaging was done on whole cleared brains from juvenile (∽P25) control and sibling *Cnr1*^*-/-*^ KO mice carrying the striosomal fluorescent marker P172-mCitrine (shown in white). ***A***, Sagittal maximum-projection images show striosomal neurons clustered in patches (examples at red asterisks) within the striatum. Striosomal axon projections are visible in the external and internal globus pallidus (arrow and arrowhead, respectively) and the rostral and caudal portions of the substantia nigra (red crosses indicate location of bouquets and red x indicates location of posterior dopamine cell cluster regions). Anterior is to the left. ***B***, Maximum-projection coronal view of the rostral nigra and all of its bouquets made visible by P172-mCitrine fluorescence in striosomal axons. In *Cnr1*^*-/-*^ KO mice compared to controls, ‘stems’ of bouquets (examples designated by red crosses) appear thinner whereas the SNcv/SNr border (red-bracketed regions) appear thicker. Medial is to the left. Scale bars are 400 μm in (***A***) and 100 μm in (***B***). See corresponding 3-D **Movies 1** and **2**.

### Striosomal and dopaminergic neuropil is disorganized at the SNcv/SNr border

To confirm that the bunched striosomal fibers in *Cnr1*^*-/-*^ KO mice were indeed located at the SNcv/SNr border, we co-immunolabeled individual brain sections for the dopamine transporter (DAT) to allow detection of dopamine-containing neurons and their processes. In control mice, the dopamine-containing cell bodies of the SNcv formed a tight layer with a sharp border between them and the SNr neuropil (**Fig. *2A***,***C***,***E***). In sharp contrast, *Cnr1*^*-/-*^ KO mice had a markedly less discrete SNcv/ SNr border and apparent disorganization of both dopaminergic dendrites (DAT-positive) and striosomal axons (P172-mCitrine-positive) (**Fig. 2*B***,***D***,***F*** and **Extended Data Fig. 2-1*A***,***B***). Although the overall distribution of dopaminergic neurons was reported to be normal in *Cnr1*^*-/-*^ KO mice (Steiner et al., 1999; Gargano et al., 2020), our new findings show that their fine-scale dendritic arbors, as well as their input circuits, were severely disrupted.

**Figure 2.**
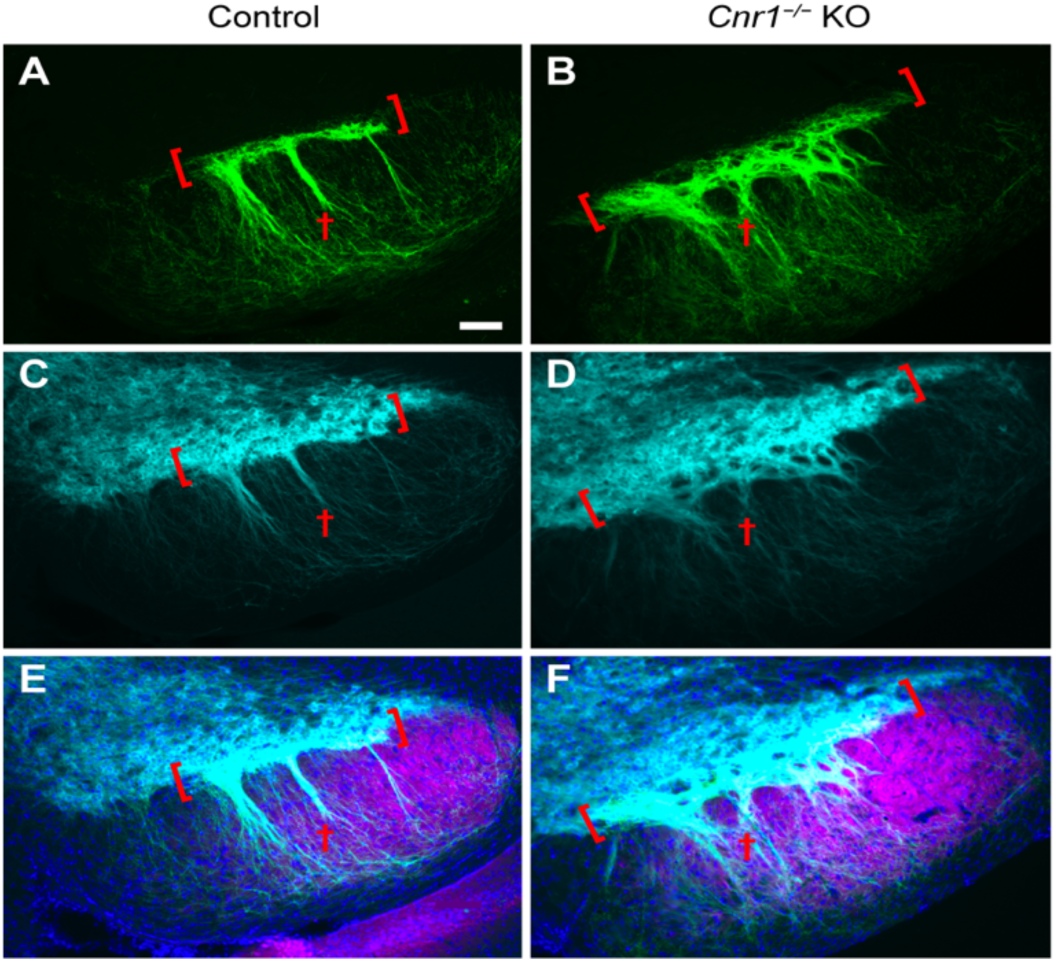
Striosomal axons and dopaminergic dendrites in the SN of *Cnr1*^*-/-*^ KO mice. Sections through the SN (left hemispheres) immunolabeled for the striosomal marker P172-mCitrine (***A, B***), the dopaminergic cell marker dopamine transporter (DAT; ***C, D***) and merged with an SNr marker in magenta (***E, F***). In *Cnr1*^*-/-*^ KO relative to control, both striosomal and DAT-positive neuropil appear unorganized and looser, failing to define either a sharp SNcv/SNr border (red brackets) or discrete bundles of striosomal axons and dopamine dendrites bundles in the SNr (red crosses). Scale bar is 100 μm and applies to all panels. N>3 for each genotype. See **Extended Data Figure 2-1 *A***,***B*** for more samples.

*P172-mCitrine* transgene expression in the striatum was previously noted to be diminished in the striatum by 5.5 weeks of age (Crittenden et al., 2016), presumably owing to developmental changes in the enhancers that control transgene expression (Shima et al., 2016). This age-dependent loss of *P172-mCitrine* expression raised the importance of using age-matched or sibling mice in order to compare expression in *Cnr1*^*-/-*^ KO mice and controls, as described in the Methods. We nevertheless capitalized on the striosome-selective loss of expression to bolster the evidence that the P172-mCitrine-labeled fibers in bouquets arise from striosomal neurons. This is particularly relevant here because cortical neurons exhibit labeling in *P172-mCitrine* transgenic mice and cortical neurons were previously shown to have axon pathfinding defects in *Cnr1*^*-/-*^ KO mice (Wu et al., 2010; Alpar et al., 2014; Saez et al., 2020). We reasoned that if there were an age-dependent loss of labeling in striosomal neurons in *P172-mCitrine* mice, then we should see a corresponding loss of fiber labeling in bouquets. Indeed, in “aged” (70 day old) *P172-mCitrine* mice we that we examined, P172-mCitrine labeling was maintained in the cerebral cortex but only a few remaining neurons remained labeled in the striosomes (**Fig. 3 *A***,***B*)**, along with only a few remaining fibers labeled in striosome-dendron bouquets. The individually labeled striosomal axons appeared to have varicosities that were in contact with bundled dopaminergic fibers descending into the SNr as well as axon end bulbs around dopaminergic fibers and neurons in the SNcv, bordering the SNr (**Fig. 4 *A-C*)**. These results support our conclusion that the disorganized bouquet fibers in *Cnr1*^*-/-*^ KO mice arise from neurons within striosomes, which are normally enriched for CB1R expression (Davis et al., 2018).

**Figure 3.**
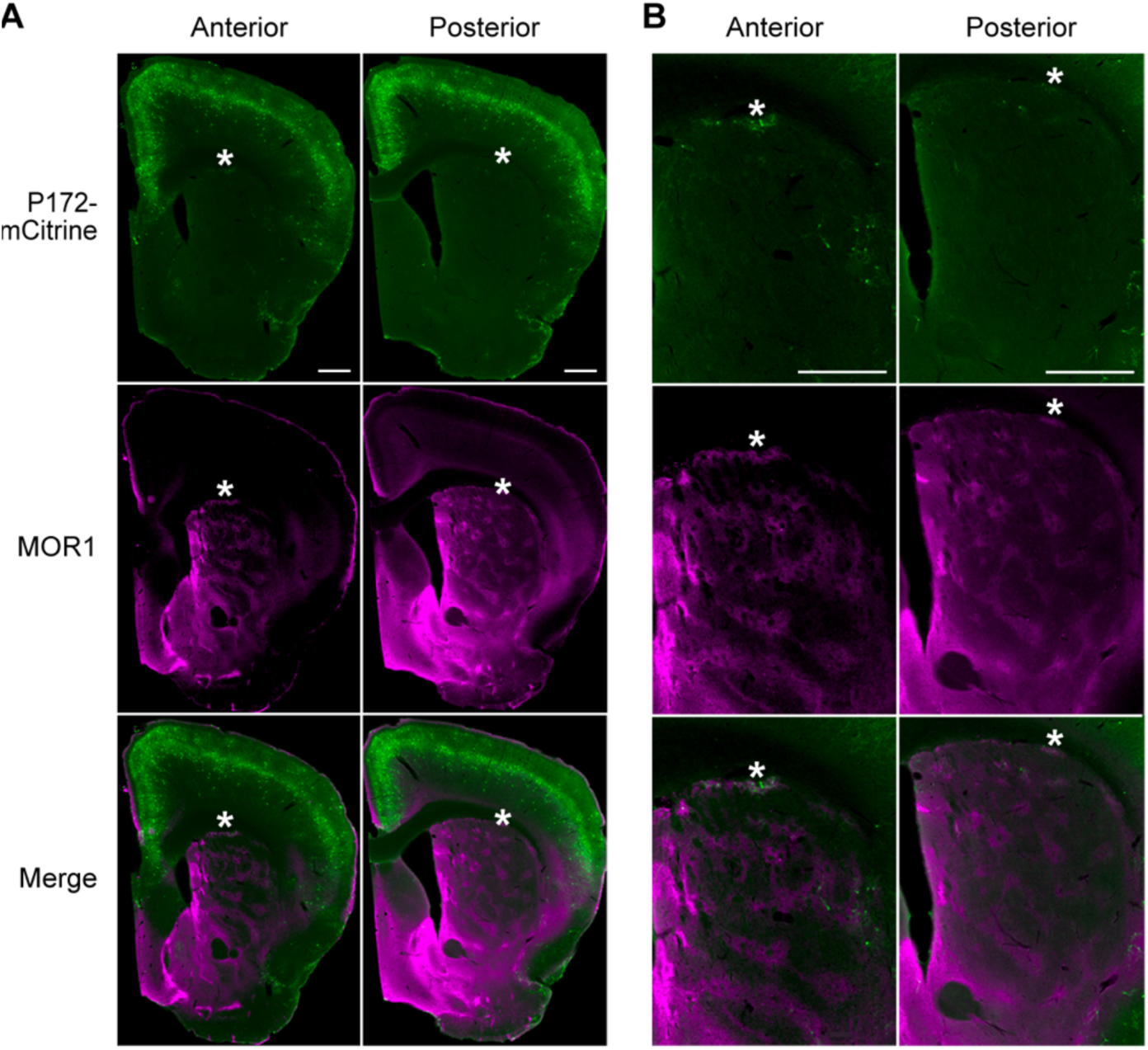
Sparse striosomal cell labeling in aged P172-mCitrine transgenic mice. ***A***, By P172-mCitrine fluorescence (green) many cells are visible in the cerebral cortex, but only a few cells remain labeled in the striosomes (identified by MOR1 labeling in magenta) of transgenic mice at 70 days of age. ***B***, Higher magnification view of sections in (***A***) showing P172-mCitrine-positive neurons in striosomes in the striatum. Coronal hemispheric sections through the anterior (top row) and posterior (bottom row) striatum are shown. Medial is to the left. Scale bars are 500 μm in top panels (***A***,***B***). Similar results in *N* = 3 mice aged 70 days. Asterisks are inserted above examples of striosomes located within the subcallosal streak.

**Figure 4.**
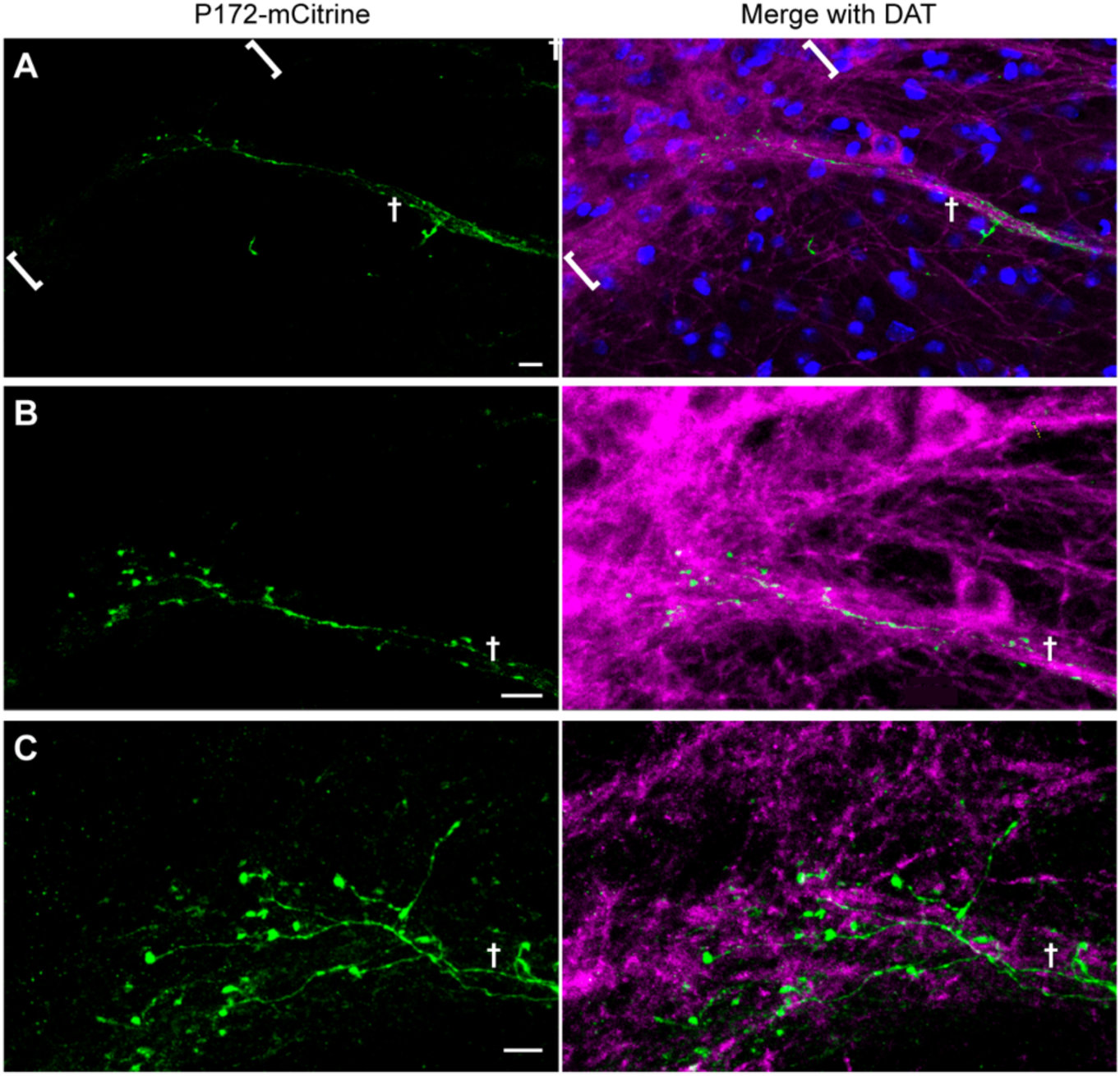
Sparse striosomal axon labeling in bouquets of aged P172-mCitrine transgenic mice. ***A-C***, Increasingly magnified images (top to bottom) show what appears to be a single P172-mCitrine (green) striosomal fiber climbing a bundle of dopaminergic dendrites labeled for DAT (magenta) and branching to form multiple end bulbs in the SNcv bordering the SNr. DAPI labels all cell nuclei in blue. The SNcv is delineated by brackets in (***A***) and the dendron is designated by a cross in (***A-C***). Scale bars are 10 μm in (***A***,***B***) and 5 μm in (***C***). *D* shows a a maximum projection through a total of 18 0.5 μm sections. Similar results in *N* = 3 mice aged 70 days. Nigral hemispheric sections are shown with medial aspect to the left.

Relative to the P172-mCitrine marker, MOR1 is stably expressed in striosomes as mice age, although it is not as strongly and discreetly expressed in striosomal bouquet axons (Crittenden et al., 2016). For genotype comparison of the bouquet phenotype in fully adult mice, we co-labeled sections for DAT and MOR1. We measured both the dorsal-ventral thickness of the SNcv/SNr border region and of the bouquet stem width in brain sections that contained bouquets from *Cnr1*^*-/-*^ KO and control mice. Bilateral measures were made by a person blind to genotype. We found that the SNcv/SNr border region, as defined by co-labeling for MOR1 and DAT, was significantly thicker in *Cnr1*^*-/-*^ KO mice than in controls (*p* = 0.0076, left hemisphere; *p* = 0.042, right hemisphere by unpaired Student’s *t*-test, *N* = 4 mice per genotype, equally divided and matched for sex, *n* = 6 hemisphere slices per mouse; **Fig. 5*A*** and **Extended Data Fig. 5-1**). This finding is consistent with what we observed by eye in single random sections and in whole-nigral movies labeled for the striosomal marker P172-mCitrine (**Fig. 1*B*, Movies 1 and 2, Fig. 2, Extended Data Fig. 2-1**). However, the widths of the bouquet stems were not significantly different between genotypes based on MOR1 and DAT labeling (*p* = 0.70 and *p* = 0.85 for MOR1 and *p* = 0.64 and *p* = 0.43 for DAT for left and right hemispheres, respectively, by unpaired Student’s *t*-test, *N* = 4 mice per genotype, equally divided and matched for sex, *n* = 6 hemisphere slices per mouse; **Fig. 5*B***,***C***). For technical reasons, we did not examine their taper.

**Figure 5.**
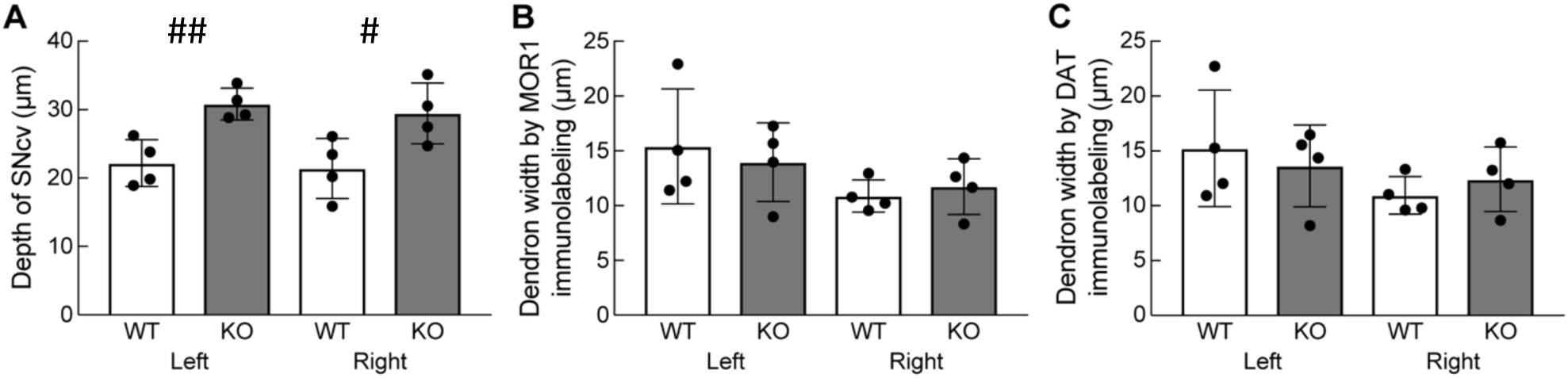
Abnormally thick SNcv in adult *Cnr1*^*-/-*^ KO mice. ***A***, The average dorsoventral depth of the SNcv border, defined as immunopositive for both MOR1 and DAT, was significantly greater as measured in coronal nigral sections from *Cnr1*^*-/-*^ KOs than in controls. (## *p* = 0.0076 for left hemisphere and ## *p* = 0.042 for right hemisphere). ***B***,***C***, The averaged mediolateral width of the dendron, measured by either MOR1 or DAT labeling at 50 μm and 100 μm ventral to the SNcv/SNr border, was not significantly different between controls and *Cnr1*^*-/-*^ KO (*p* = 0.70 and *p* = 0.85 for MOR1 and *p* = 0.64 and *p* = 0.43 for DAT for left and right hemispheres, respectively). Left and Right designate opposite hemispheres from the same mice. Means for each animal and standard deviations of interanimal variability are plotted. Images from which measurements were made are shown in **Extended Data Figure 5-1**.

### There were few significant differences in MOR1 and DAT immunolabeling in *Cnr1*^*-/-*^ KO mice vs. controls

We also compared the MOR1 and DAT markers in *Cnr1*^*-/-*^ KO mice and controls according to their immunointensity. We measured MOR1 and DAT intensity within the SNcv and within dendrons in the regions used for measuring SNcv thickness and dendron width. No significant genotype differences were found (by unpaired Student’s *t*-test, *N* = 4 mice per genotype, equally divided and matched for sex, *n* = 3 left hemisphere slices per mouse, *p* > 0.05 for all comparisons between genotypes except for MOR1 immunointensity in left hemisphere dendrons for which *p* = 0.046; **Extended Data 5-2*A-D***). The trend for less MOR1 and DAT immunolabeling of dendrons in the left hemisphere of *Cnr1*^*-/-*^ KO mice (**Extended Data 5-2*C***,***D***) could be related to the tendency for sparser dendrons as seen in some samples (**Fig. 1*B***, **Movie 2**, **Fig. 2*B***,***D***,***F*, Extended Data Fig. 2-1*B***). The finding that DAT and MOR1 immunointensity in the SNcv were not different from controls is important to support that the thickened SNcv phenotype in *Cnr1*^*-/-*^ KO mice is not an artifact of stronger fiber labeling.

Tests for changes in MOR1 expression in the striatum are particularly relevant for *Cnr1*^*-/-*^ KO mice, which we report here have anatomical abnormalities in their SNcv dopamine systems, because of previous findings indicating (1) that disruption of SNcv dopaminergic neuropil leads to reduced MOR1 immunolabeling in striata of rodents (Johansson et al. 2001) as a possible compensatory response to increased levels of the MOR ligand enkephalin detected by immunolabeling (Koizumi et al., 2013) and (2) that enkephalin itself is increased in the striata of *Cnr1*^*-/-*^ KO mice (Steiner et al., 1999). We measured MOR1 immunoreactivity within striosome and matrix compartments of the striatum at three anterior-posterior levels (**Fig. 6*A, B***) but found no genotype differences between *Cnr1*^*-/-*^ KO mice and controls (*p* > 0.05 for all comparisons between genotypes by unpaired Student’s *t*-test, *N* = 4 mice per genotype, equally divided and matched for sex, *n* = 3 hemisphere slices per mouse at matched levels; **Fig. 6*C, D***).

**Figure 6.**
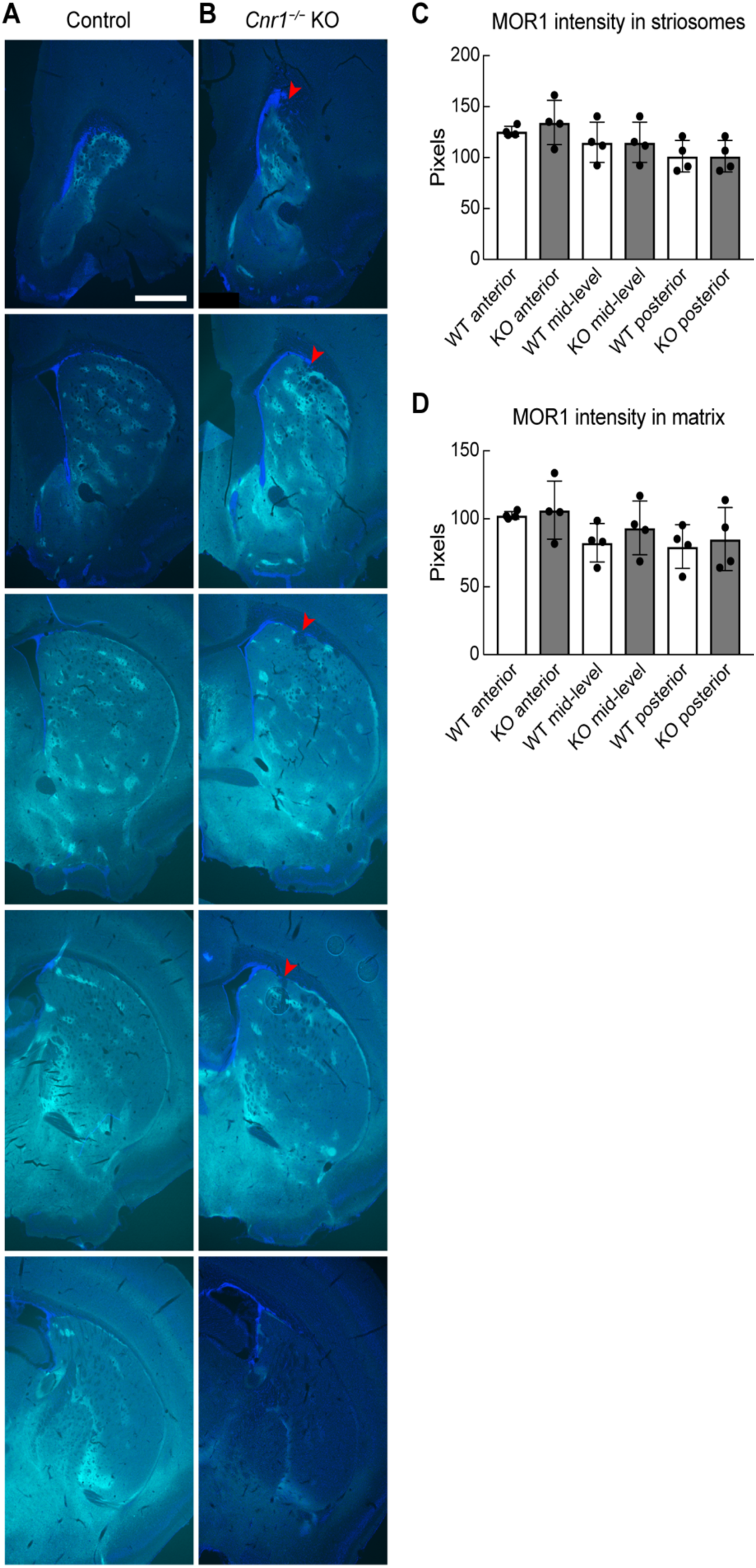
MOR1 striosomal labeling in adult *Cnr1*^*-/-*^ KO mice. ***A***,***B***, The expression level and pattern of labeling for the striosomal immunomarker MOR1 (cyan) are similar in serial sections from a control (***A***) and a *Cnr1*^*-/-*^ KO (***B***) mouse. The left striatal hemisphere is shown from anterior (top) to posterior (bottom). Cell nuclei are labeled by DAPI (blue). White arrowhead in *Cnr1*^*-/-*^ KO panels indicates the abnormally bunched corticofugal fibers near the border of the dorsolateral striatum. Scale bar in the upper left panel is 500 μm and applies to all panels. *N* = 5 mice imaged for each genotype. ***C***,***D***, Average MOR1 immunointensities (arbitrary units) in striosomes and matrix compared between genotypes at three anteroposterior levels. The middle three sections shown in (***A***,***D***) are representative of the levels chosen. No significant genotype differences were found (*p* > 0.05 for all comparisons). Means for each animal and standard deviations of interanimal variability are plotted.

### Striosomal areas and neuronal counts were not significantly different in *Cnr1*^*-/-*^ KO mice vs. controls

To determine whether the terminal field abnormalities of the striosome-nigral axons reflected a distortion of striatal striosome organization, we examined striatal anatomy with the striosomal neuropil marker MOR1 and the striosomal cell body markers P172-mCitrine and FoxP2. Our finding that MOR1 immunolabeling intensity in the striatum of *Cnr1*^*-/-*^ KO mice was not significantly different from controls (**Fig. 6*A-D***) enabled us to use MOR1 as a marker to identify striosomal and matrix areas in all of these assays.

In *Cnr1*^*-/-*^ KO mice, we observed that MOR1-positive clusters were perturbed at the dorsolateral aspect of the caudoputamen, in the region where abnormally bunched corticofugal fibers enter the striatum, as previously described for *Cnr1*^*-/-*^ KO mice (Alpar et al., 2014; Berghuis et al., 2007; Mulder et al., 2008; Saez et al., 2020; Wu et al., 2010; **Fig. 6*B***). However, we found no significant differences in the total striatal, striosomal and matrix areas and the striosome size and counts per section between *Cnr1*^*-/-*^ KO and control mice when matched levels were compared (*p* > 0.05 for all genotype comparisons by Student’s *t*-test, N = 4 mice per genotype, *n* = 3 matched-level slices per mouse; **Fig. 7*A-D***). As expected, the average MOR1-labeled areas were larger on average in the anterior part than in the posterior part of the striatum (**Fig. 7*B***,***D***; Miyamoto et al., 2018).

**Figure 7.**
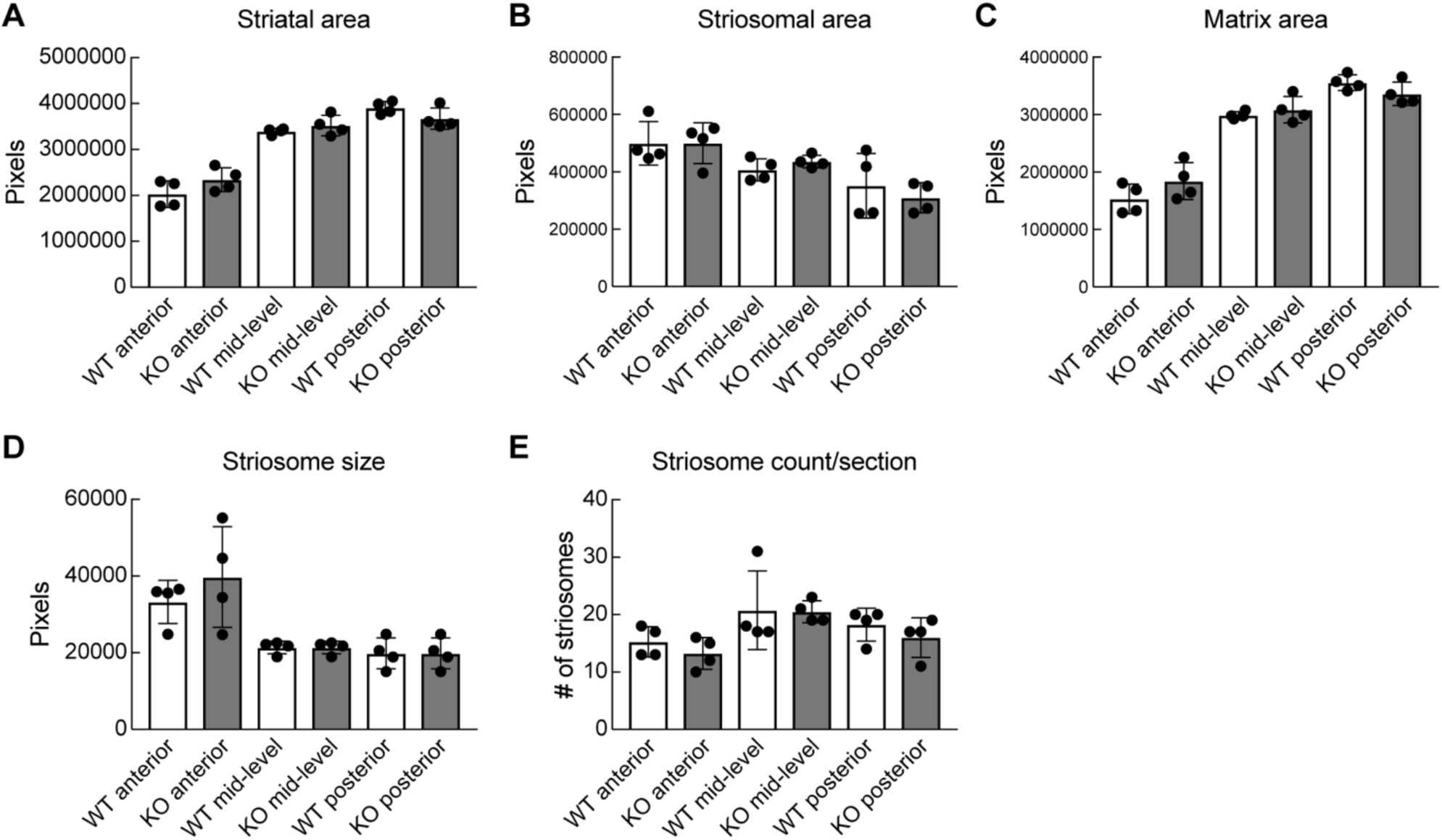
Striosome-matrix architecture in adult in *Cnr1*^*-/-*^ KO mice. MOR1 immunolabeling in coronal sections at matched levels were used to test for genotype differences in the total area of the striatum (***A***), striosomes (***B***) and matrix (***C***) and for measuring average striosome size (***D***) and density (***E***). No significant genotype differences were found (*p* > 0.05 for all comparisons). Means for each animal and standard deviations of interanimal variability are plotted.

As a further test of striatal compartmentalization, we counted the number of striosomal cell bodies that were positive for P172-mCitrine-or FoxP2-immunolabeling in sections from adult *Cnr1*^*-/-*^ KO, *Cnr1*^*−/+*^ heterozygous and sibling control mice. We found that the densities within MOR1-positive striosomes and across the whole striatum were not significantly different between genotypes (**Fig. 8*A-J***). These counts of putative striosomal neurons still do not address subtle changes in the disposition of the striosomes as seen in cross-section or as seen in 3D in *Cnr1*^*-/-*^ KO mice, particularly at the dorsolateral border with the white matter, which might be related to the abnormal patterning of cerebral cortical and thalamic axons that course through the striatum. Our findings nevertheless indicate that CB1R signaling is neither required for the ultimate formation of striosomal neurons, nor for their organization into clusters (Matsushima & Graybiel, 2020). This situation stands in contrast to the well documented requirement for CB1R in cell neogenesis, survival or differentiation in other brain regions including the cerebral cortex (Gaffuri et al., 2012), a phenotype evidenced by the paucity of P172-mCitrine-positive cells that we observed in cortical layer 4 (Shima et al., 2016) of *Cnr1*^*-/-*^ KO mice (**Fig. 8*C, D***).

**Figure 8.**
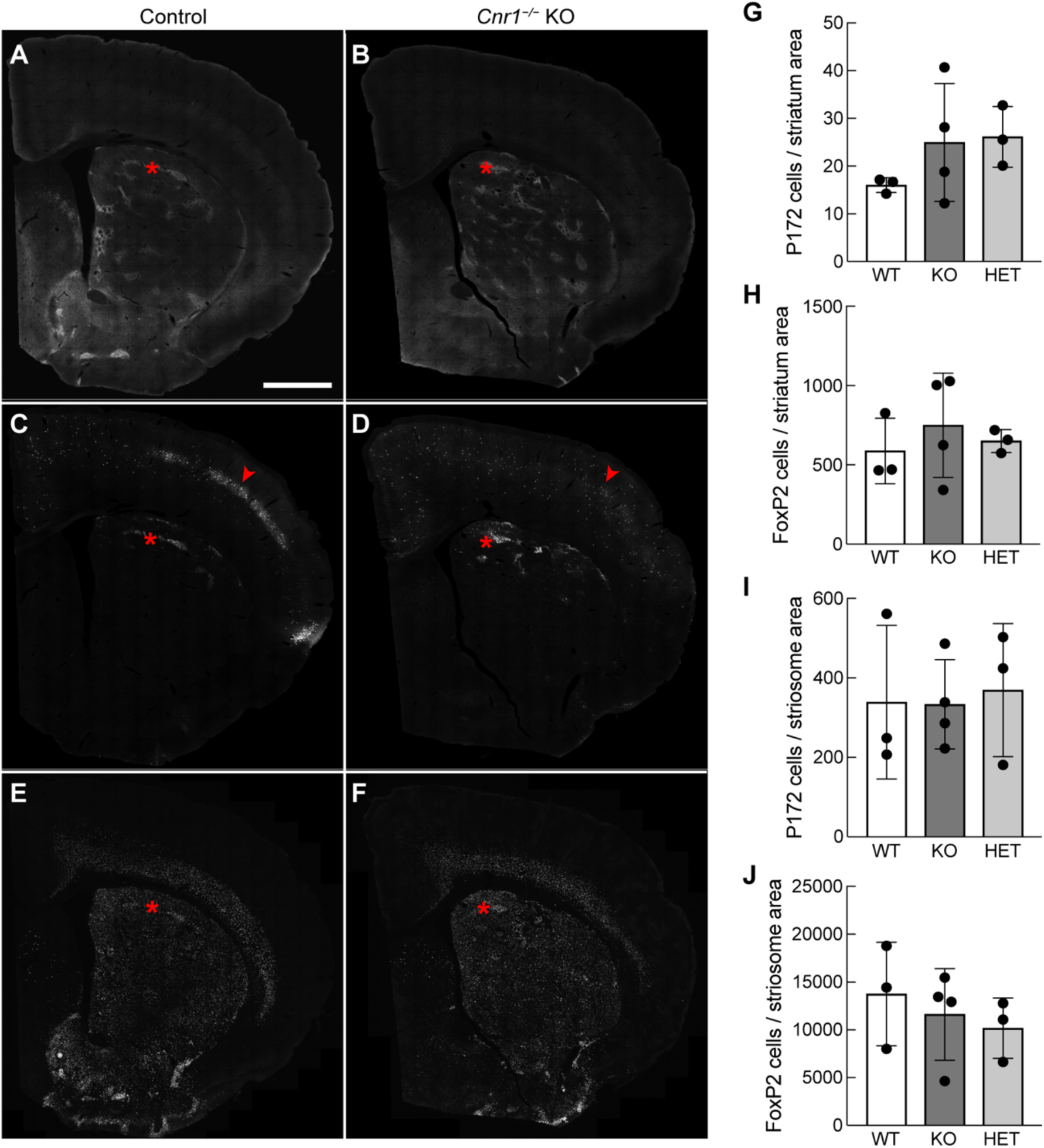
Striosomal cell density in adult in *Cnr1*^*-/-*^ KO mice. Samples of striatal sections triple-labeled for MOR1 (***A***,***B***), *P172-mCitrine* (***C***,***D***) and FoxP2 (***E***,***F***) that were used for counting striosomal cell densities within striosomes and across the dorsal striatum. Red asterisks highlight single striosomes. Note the absence of a *P172-mCitrine*-labeled cells in cortical layer 4 in *Cnr1*^*-/-*^ KO mice (region designated by a red arrowheads in control (***C***) and KO (***D***)). Scale bar in (***A***) is 500 μm and applies to all panels. Graphs show the number of cells labeled for the striosomal cell markers P172-mCitrine (***G, I***) and FoxP2 (***H, J***), normalized to striatal (***G***,***H***) or striosomal (***I***,***J***) area (arbitrary units). Cells were counted for each of 4 level-matched sections from control (*N* = 3), *Cnr1*^*-/-*^ KO (*N* = 4) and *Cnr1*^*−/+*^ HET (*N* = 3) mice. For all comparisons to controls, p > 0.05 by ANOVA with Dunnett’s correction. Means for each animal and standard deviations of interanimal variability are plotted.

### In *Cnr1*^*-/-*^ KO mice, striosomal and dopamine-containing neurons successfully reached their long-range targets via the striatonigral and nigrostriatal tracts

The dopamine-containing neurons of the ventral tier SNc, located within the poorly organized striosomal neuropil in *Cnr1*^*-/-*^ KO mice, are known to send long-range axonal projections to the dorsal striatum, with a preference for striosomes (Matsuda et al., 2009; Crittenden et al., 2016; Sgobio et al., 2017). A smaller proportion of dopamine-containing inputs to the dorsal striatum arise from the ventral tegmental area of the midbrain, which was not studied here. In adult *Cnr1*^*-/-*^ KO mice, the nigrostriatal projections appeared generally normal, based on immunolabeling for the dopamine transporter (**Fig. 9*A-C***). In newborn mice, dopamine-containing inputs preferentially target developing striosomes to form tyrosine hydroxylase (TH)-positive “dopamine islands” (Graybiel, 1984; Miura et al., 2007). We did not perform quantitative analyses, but we did observe typical distributions of TH-positive dopamine islands that corresponded to striosomes based on overlapping distribution of P172-mCitrine-positive cells in postnatal day (P) 5 control and *Cnr1*^*-/-*^ KO brains (**Fig. 10*A-D***). We detected striosome-nigral development with P172-mCitrine and MOR1 immunomarkers for striosomal axons and TH for dopaminergic dendrites; DAT expression was not used because it was not enriched in dopaminergic nigral processes of the early postnatal mice. It was clear that burgeoning bouquets were present in the SN with both striosomal axon and dopaminergic dendrite markers, in P5 *Cnr1*^*-/-*^ KO pups (**Fig. 10*E, F***).

**Figure 9.**
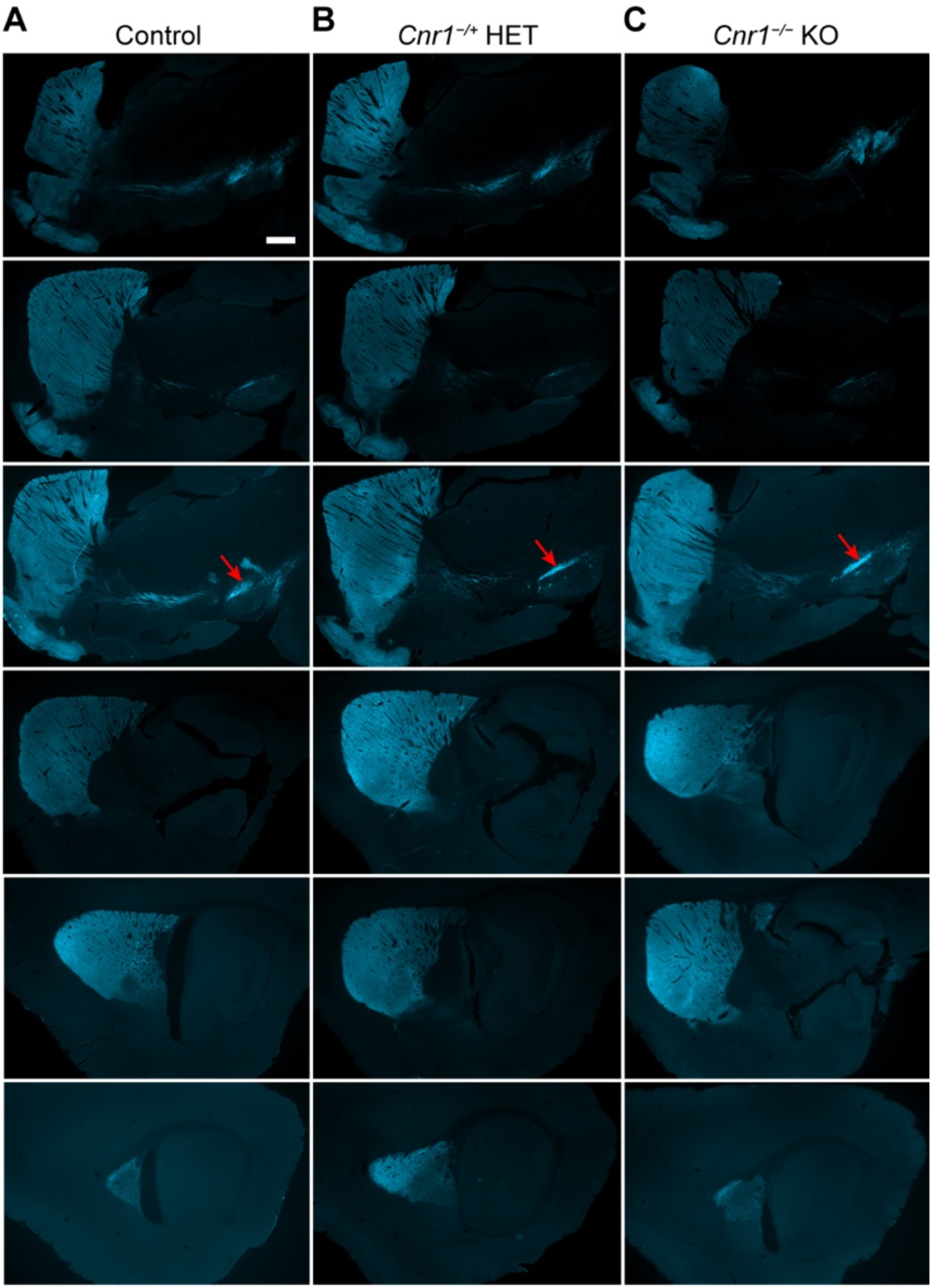
Nigrostriatal innervation in adult *Cnr1*^*-/-*^ KO mice. Serial sagittal sections through the basal ganglia of control (***A***), *Cnr1*^*−/+*^ heterozygous (***B***) and *Cnr1*^*-/-*^ KO (***C***) mice immunolabeled for dopamine transporter DAT (cyan) highlight the dopaminergic projections from the midbrain that innervate the entire striatum. Anterior is to the left and sections run from medial (top) to lateral (bottom). The ventral tier dopaminergic neurons are indicated by a red arrow. Scale bar in upper left panel is 500 μm and applies to all panels. *N* = 5 mice imaged for each genotype.

**Figure 10.**
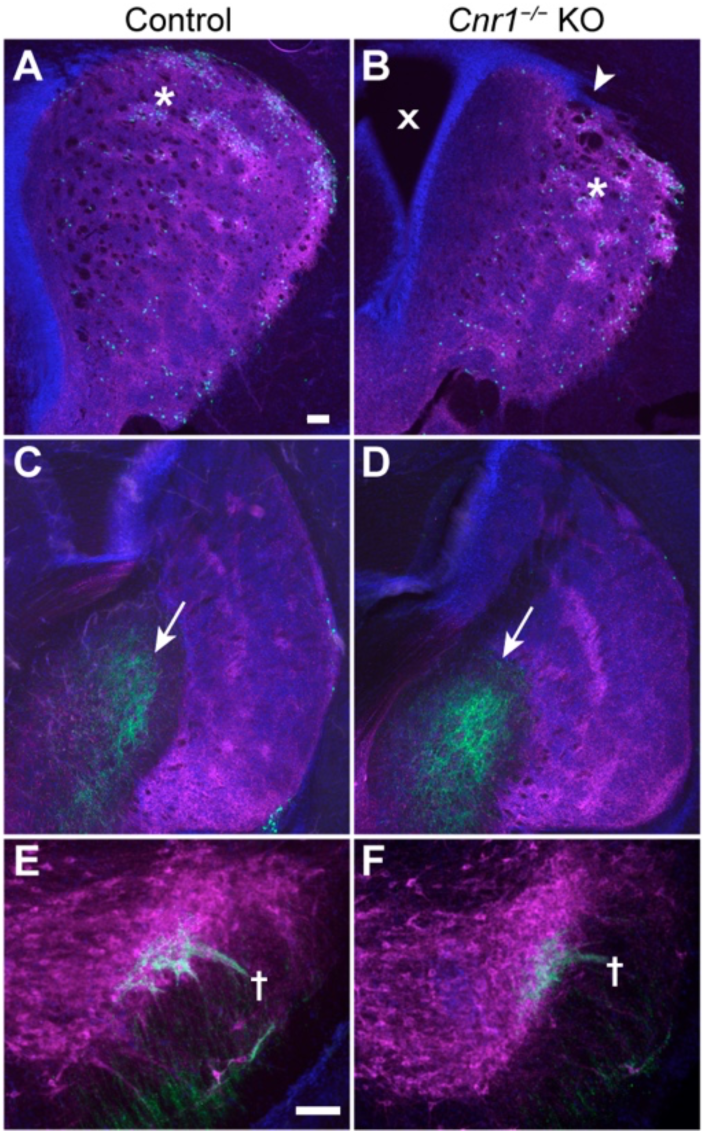
Developing striosomes and dopamine islands in *Cnr1*^*-/-*^ KO mouse pups. Serial coronal sections through the basal ganglia of P5 pups carrying the P172-mCitrine striosomal marker (green) are immunolabeled for the dopaminergic cell marker TH (magenta) of controls and *Cnr1*^*-/-*^ KO pups. Medial is to the left, lateral to the right for each panel. Dopamine islands (white asterisks in top panels) corresponding to developing striosomes are enriched for TH fibers and P172-mCitrine-labeled (mostly dorsal) striosomal neurons in the anterior (***A***,***B***) and posterior (***C***,***D***) striatum. A white arrowhead (***B***) indicates where the striatum is disrupted by bunched corticofugal fibers piercing the dorsolateral striatum, a known *Cnr1*^*-/-*^ KO phenotype (Wu et al., 2010). The enlarged ventricles (marked by an x in (***B***)) and laterally confined striosomes and dopamine islands suggest delayed striatal development in some *Cnr1*^*-/-*^ KO pups relative to sibling controls but this was not further investigated. Striosomal fibers reaching the external globus pallidus (marked by arrows in (**C**) and (**D**)) are evident in both genotypes by P172-mCitrine labeling (***C***,***D***). ***E***,***F***, The proximity of striosomal axons and dopaminergic fibers in bouquets is evident in controls (unresolved green and magenta labeling appears white) but in *Cnr1*^*-/-*^ KO pups the co-labeled bundles are less distinct (developing bouquet stems are indicated by crosses). Scale bars in (***A***,***E***) are 100 μm and apply to all panels. *N* = 4 pups examined for each genotype. Cell nuclei are labeled by DAPI (blue).

### Striosome-dendron bouquets formed, and became enriched for CB1R, between postnatal days 5 and 7

To test for a potential influence of CB1R expression on the initial formation of bouquets, we determined when striosomes and bouquets were forming in developing control pups and when CB1R expression began in incoming striosomal axons destined to innervate the bouquet dendrons. In P0 to P11 pups, we found that we could track the initial formation and progressive formation of striosomes and growth of bouquet stems (**Fig. 11*A, B***). Remarkably expression of CB1R in striosomes and striosome-nigral axons (labeled for P172-mCitrine) was not yet visible at P5 (**Fig. 11*A, B***), despite the abundance of CB1R in corticofugal fibers from P0 to P5 (Harkany et al., 2007). By P7 however, CB1R expression in corticofugal fibers passing through the striatum was diminished and CB1R expression in striatal neuropil, with enriched expression in striosomes, became apparent (**Fig. 11*A***). Correspondingly enrichment for CB1R within striosomal axons in bouquet stems and at the SNcv/SNr border, relative to the surrounding nigral neuropil, also became apparent by P7 (**Fig. 11*B***).

**Figure 11.**
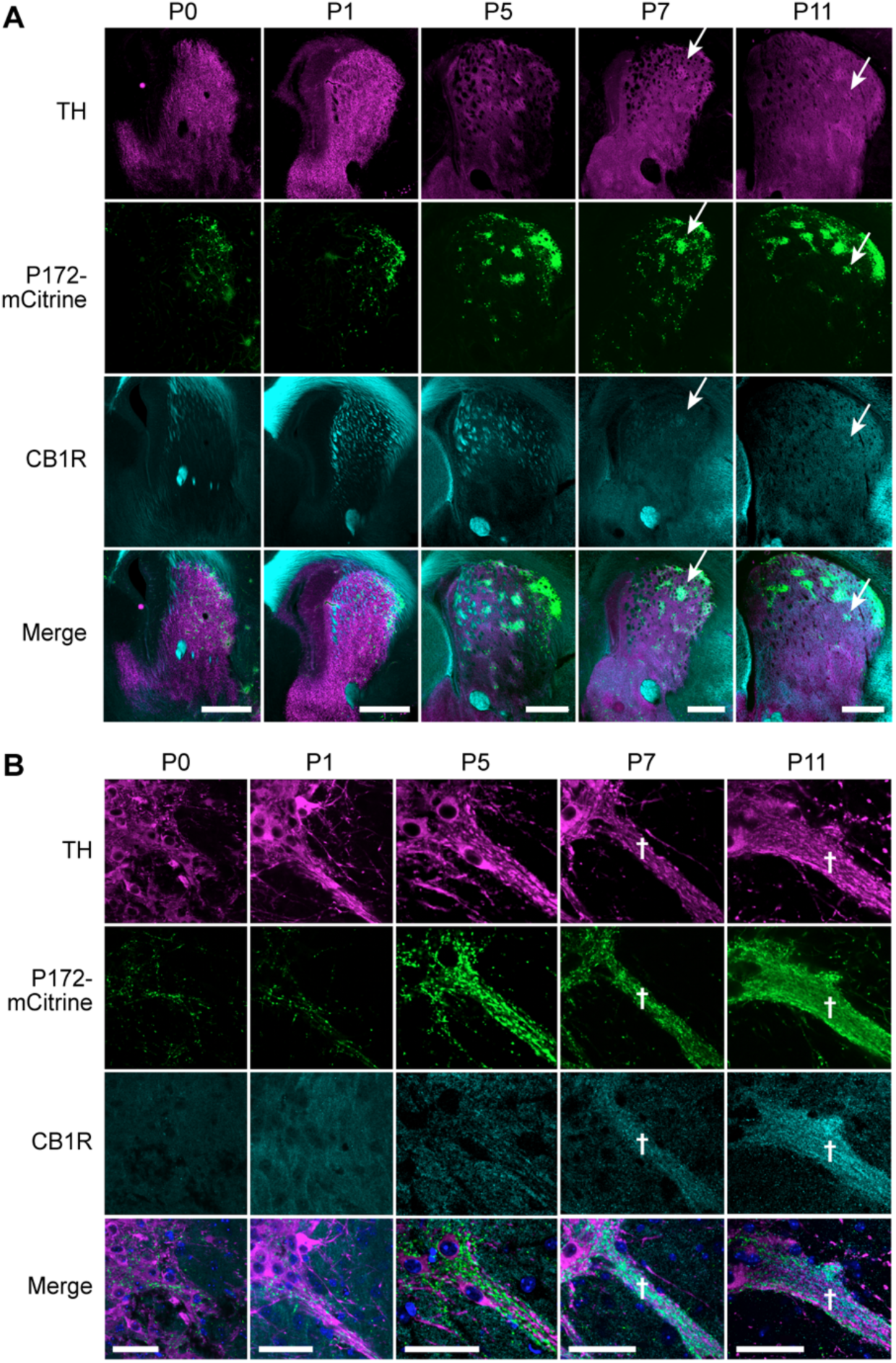
Bouquet formation in neonatal mice coincides with CB1R enrichment in striosomes. ***A***, Coronal sections through the left striatum in P0-P11 mouse pups, co-labeled to identify dopamine islands that innervate developing striosomes (TH) and striosomal markers (P172-mCitrine and CB1R). CB1R enrichment in striosomes is first apparent in the sections from P7 pups (white arrows in P7 and P11 panels), which corresponds to its enrichment in the expanding bouquet ‘stem’ (designated by white asterisks in panel (***B***)). A white arrow designates a single striosome labeled for TH, P172 and CB1R in P7 and P11 pup brains. Scale bars are 500 μm and shown in the merged images panel. N = 4 pups for each age. ***B***, Coronal sections through the left substantia nigra show single bouquets, from P0-P11 mouse pups, co-labeled to identify dopaminergic dendrites (TH), and striosome axons (P172-mCitrine and CB1R). CB1R expression onset in striosomal fibers near the TH-rich SNcv border and along ventrally-extending TH-positive fiber bundles appears to correlate with the enlargement of the bouquet ‘stem’ (defined by white crosses in P7 and P11 panels). Scale bars are 50 μm and shown in the merged images panel. *N* = 4 pups for each age.

We considered that the gradual postnatal enrichment of CB1R in developing bouquets might simply be a function striosomal axon density, rather than rising expression in each fiber. However, the expression of P172-mCitrine was strongly expressed in bouquet already by P5, and the gradual striosomal enrichment of CB1R expression at the level of the striatum during this early postnatal period (**Fig. 11*A***) further indicated that it was not until after P5 that CB1R became enriched in striosomal neurons and their nigral projections. A key control supporting this conclusion was confirmation of the specificity of the CB1R antibody that we used. We found strong CB1R antibody immunoreactivity in adult controls, with slightly reduced immunoreactivity in *Cnr1*^*−/+*^ heterozygous mice and no labeling in *Cnr1*^*-/-*^ KO mice (**Fig. 12**).

**Figure 12.**
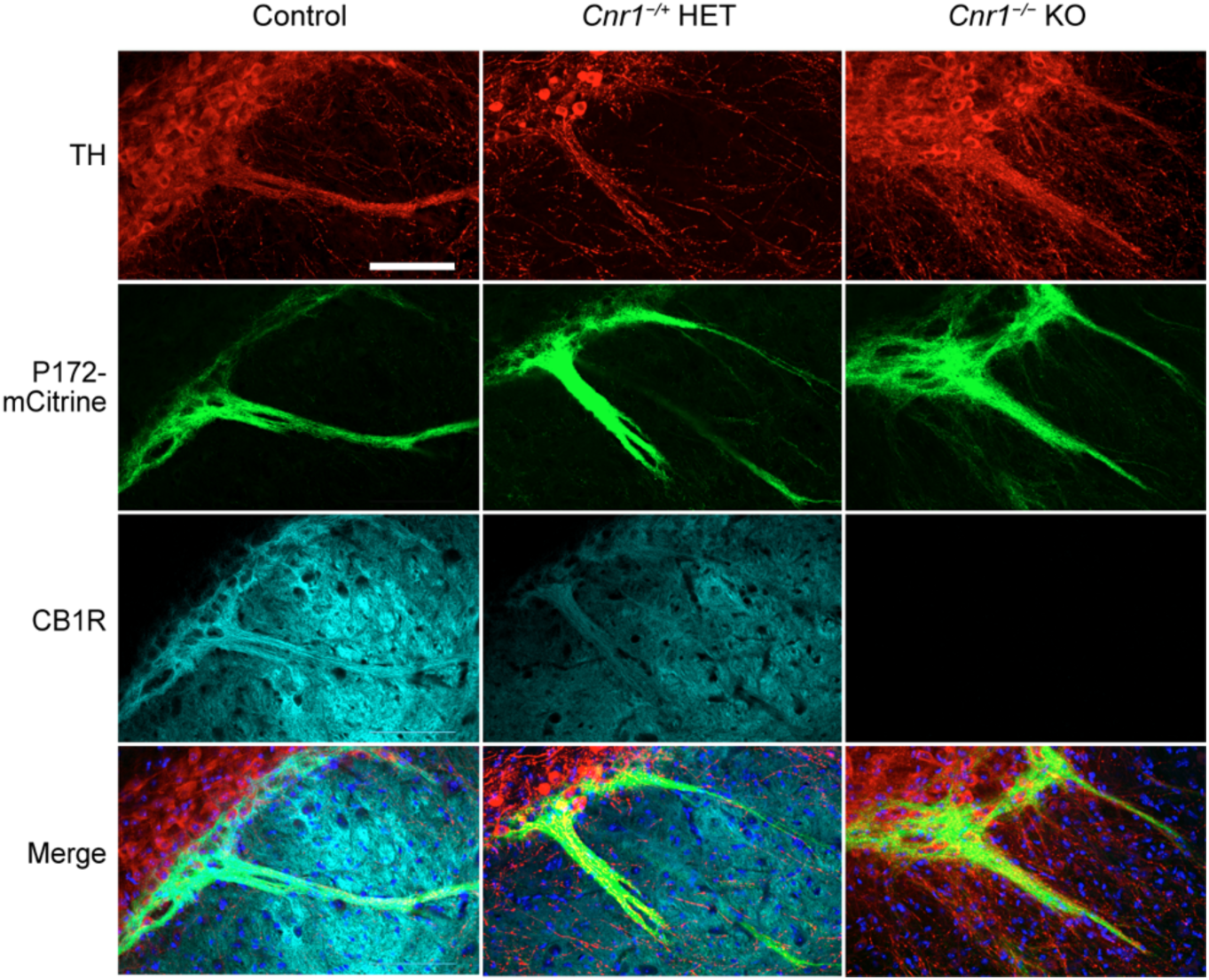
CB1R antibody specificity. Bouquets of adult *Cnr* WT, HET and KO mice carrying *P172-mCitrine* were immunolabeled for CB1R and TH, confirming immunospecificity of the CB1R antibody. N > 3 mice per genptype. Scale bar is 100 μm and applies to all panels.

Finally, we tested whether the bouquet phenotype in adult *Cnr1*^*-/-*^ KO mice was already evident at ages corresponding to the onset of CB1R enrichment in striosomal axons. It was: in P11 pups the striosomal axons and dopamine-containing dendrites clearly appeared piled up at the SNcv/SNr border (**Fig. 13*A, B*)**. We quantified this phenotype by measuring, genoty#pe blind, the dorsal-ventral distance of the region co-labeled for MOR1 and TH and found that there was a significantly thicker SNcv/SNr border region in P11 *Cnr1*^*-/-*^ KO mice than in controls (*p* = 0.029 by unpaired Student’s *t*-test, *N* = 4 mice per genotype, *n* = 1 hemisphere slice per mouse pup; **Fig. 13*C***). All together, these results suggest that CB1R expression begins in striosomal axons postnatally, during bouquet formation, and is required for striosomal axons and their terminals to communicate normally with dopamine-containing dendrites by forming discrete bouquets.

**Figure 13.**
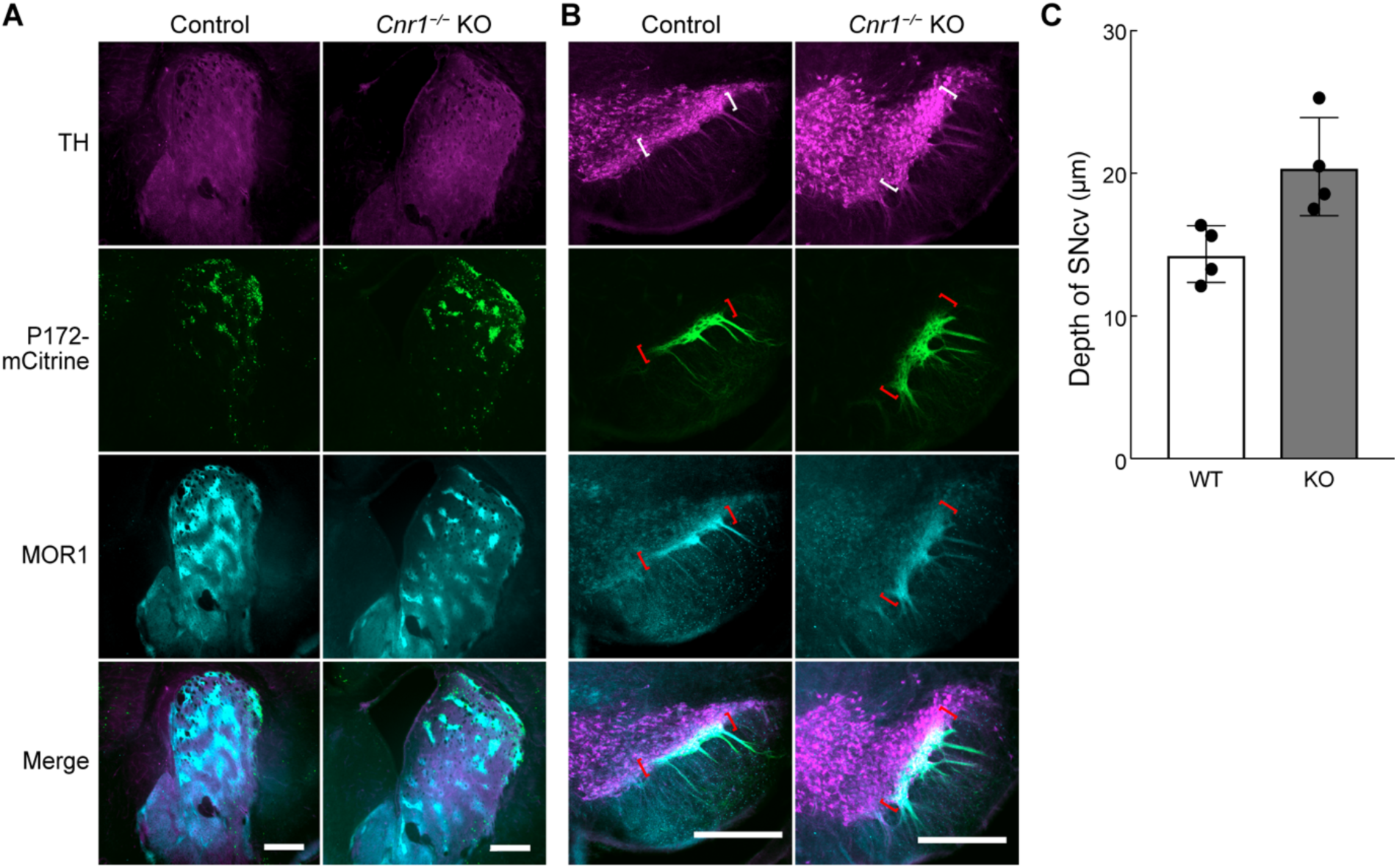
Developing bouquets are abnormal in *Cnr1*^*-/-*^ KO developing mice. Coronal hemi-sections through the left striatum ***A***, and substantia nigra ***B***, of P11 pups carrying the P172-mCitrine striosomal marker (green) are immunolabeled for the dopaminergic cell marker TH (magenta) and striosomal projection neuron marker MOR1 (cyan). Similar to the adult, the striosome organization appears grossly normal in *Cnr1*^*-/-*^ KO mice at this intermediate stage of bouquet development, but the striosomal and dopaminergic fibers appear bunched up rather than forming a discrete border (region in colored brackets) between the SNcv and the SNr as they do in control mice. *N* = 4 mice for each genotype. **C**, The average dorsoventral depth of the SNcv border, defined as immunopositive for both MOR1 and TH, was significantly greater as measured in coronal nigral sections from *Cnr1*^*-/-*^ KO pups compared to controls (# *p = 0*.*029* by Student’s unpaired *t*-test, *N* = 4 mouse hemispheres per genotype). Scale bars are 500 μm and shown in the merged images panel. Means for each animal and standard deviations of interanimal variability are plotted. Images from which measurements were made are shown in **Extended Data Figure 13-1 *A***,***B***.

## Discussion

### CB1R in neurodevelopment of striosome-dendron bouquets

Our findings lead to two main conclusions. First, it is in the early postnatal period that the elaboration of striosome-dendron bouquets occurs in the substantia nigra. These consist of clusters of dopamine-containing nigral neurons, located within the SNcv along the border with the SNr, that intertwine their proximal and ventrally extending dendrites with one another and with incoming striatal afferent fibers from striosomes. This close intertwining of striosomal inputs allows powerful control of the ventral nigral tier by clusters of striosomal neurons in the striatum. Thus, this critical early postnatal period is one of growth, but also a time of vulnerability, of the striatonigral circuit. Our second conclusion, based on observations in *Cnr1*^*-/-*^ KO mice lacking expression of CB1R, is that the endocannabinoid receptor CB1R exerts a critical and essential influence on this postnatal development of striosome-dendron bouquets. The incoming striosomal axons and the ventral dopamine-containing neurons fail to form fully organized bouquets, resulting in a pileup of incoming striosomal axons at the ventral tier border with the SNr. This finding serves as an alert that interfering with CB1R signaling, including here the *Cnr1* genetic deletion mimicking CB1R antagonism, could distort dopamine signaling beginning in the first days of postnatal development and into adulthood.

The ventral tier nigrostriatal dopamine-producing neurons are thought to modulate distinct aspects of motor control and behavior both by means of dopamine release in the dorsal striatum and by local dopamine release within the substantia nigra, including within the SNr that lies ventral to the SNcv and is penetrated by descending dopaminergic dendrites (Mukhida et al., 2001; Witkovsky et al., 2009; Zhou et al., 2009; Rice & Patel, 2015; Robinson et al., 2019; Salvatore et al., 2019; González-Rodríguez et al., 2021). Thus, disruption of striatonigral CB1R signaling in newborns could have profound long-term effects on dopamine-dependent behaviors; these could include not only movement-related activity but also aspects of mood, motivation and possibly habit formation including habitual drug abuse. These are important issues to address. Whether exogenous over-activation as well as under-activation of CB1R has neurodevelopmental consequences for the striatonigral system has not been addressed by our findings, but both conditions are of great public health consequence because cannabinoids, such as those in marijuana, are readily transmitted to nursing infants via mother’s milk and treatment with either agonists or antagonists of CB1R could possibly affect dopamine-dependent modulation of behavior and neural circuit function (Scheyer et al., 2020; Schneider, 2009).

Despite the deformations in the organization of striosomal axons and dopamine-containing dendrites within the substantia nigra of *Cnr1*^*-/-*^ KO mice, we did not observe gross changes in the density of striosomal fibers within the striatonigral tract that extends to this midbrain target. Nor did we observe gross changes in the density of dopaminergic axons in the nigrostriatal tract reaching the striatum. These findings are consistent with there not being significant changes between *Cnr1*^*-/-*^ KOs and controls in the number of striosomal neurons, based on our area measurements and cell body counts with the striosome markers MOR1, P172-mCitrine and FoxP2 or, in previous work, the number of TH-positive dopaminergic neurons in the substantia nigra (Steiner et al., 1999; Gargano et al., 2020). However, we did see deformation of the dorsolateral aspect of the caudoputamen, in regions where at minimum the striosomal so-called streak at the edge of the striatum could have been affected. Given that striosomal axons did reach the substantia nigra, even though failing to form normal bouquets, strongly suggests that CB1R function is required for the normal juxtaposition of striosomal and dopaminergic fibers within the midbrain, but that it is not essential for their long-range pathfinding. This conclusion is striking, given evidence from study of the birthdate-dependent development of the striosomal innervation of the bouquets that the long-distance trajectory and final arborization patterns of the striosome-nigral innervations are under separate control (Matsushima & Graybiel, 2020).

### CB1R in prenatal and postnatal axon pathfinding

During development of the cerebral cortex and hippocampus in rodents, CB1R is less strongly expressed in the postnatal period than it is prenatally, when it has key functions in neuronal proliferation, migration and axon pathfinding for neurons and interneurons. Long-range guidance of cerebral corticofugal axons during embryonic development (Alpar et al., 2014; Saez et al., 2020) is thought to involve a meeting known as a ‘handshake’ mechanism that occurs near the border of the striatum and the corpus callosum (Wu et al., 2010). When CB1R-expressing corticofugal axons extend ventrally into the striatum, they encounter endocannabinoid-releasing thalamocortical axons that are themselves extending dorsally into the striatum. CB1R-dependent endocannabinoid signaling between the two axonal types has been reported to allow the developing fiber connections to find their respective targets (Wu et al., 2010). In *Cnr1*^*-/-*^ KO adult and P5 mice, the cortical fibers fail to find their targets and appear bunched up near the border between striatum and corpus callosum as previously shown and as seen in **Figs. 6*B* and 10*B***.

Whether the dendrites of the dopamine-containing neurons of the developing bouquet stems produce or release endocannabinoids is unknown. However, at maturity, dopamine-containing neurons in the nearby ventral tegmental area express endocannabinoids that signal to CB1R on input axons from the nucleus accumbens septi (Riegel & Lupica, 2004). Based on this biologic principal, we hypothesize that, during development, CB1R-positive striosomal axons “handshake” with endocannabinoid-producing dendrites from dopaminergic SNcv neurons to guide the tight intertwining of striosomal axons and dopaminergic dendrites that form the normally discrete border between the SNcv and SNr and the ventrally extending bouquet dendrons. Our data in mice suggest that this signaling handshake occurs postnatally, as CB1R expression becomes enriched in striosomes and their striatonigral axons between P5 and P7, and loss of CB1R gives rise to measurably abnormal bouquets by P11. Our measurements indicated that in *Cnr1*^*-/-*^ KO adult and P11 mice, the intertwined striosomal and dopaminergic fibers formed a thicker SNcv tier, and they appeared less discrete and more loosely bundled along the border with the SNr. Expression of the striosomal axon marker MOR1 was significantly lower in bouquet dendrons in the left hemisphere, with a similar trend for the right, in adult *Cnr1*^*-/-*^ KOs relative to controls, consistent with the observed sparsity of dendrons in some *Cnr1*^*-/-*^ KO mice. All together, these results suggest that, in the absence of CB1R, striosomal axons reach the midbrain but become bunched along the SNcv/SNr border rather than organizing with dopaminergic fibers to form discrete bouquets and ventrally-extending dendrons.

By tracing of single fibers in aged *P172-mCitrine* mice, we observed one that appeared to climb dorsally along the stem and to form terminal branches and bulbs along the SNcv/SNr border. Whether all striosomal fibers follow the same path to form bouquets remains unknown. In the absence of CB1R, signaling between striosomal axons and dopamine-containing dendrites could fail, and normal connections could be disrupted as a consequence, potentially contributing to the disorganized SNcv in *Cnr1*^*-/-*^ KO mice. We noted that it was along architectural borders, both for the striatum and the nigra, that the loss of CB1R expression had the most obvious impact.

An alternative accounting for the abnormal striosome-dendron bouquet phenotype in the *Cnr1*^*-/-*^ KO mice is that the there is a temporal mismatch of the connections. For example, if striatal projection neurons undergo delayed maturation, they might not reach their normal nigral destination during the appropriate time-window to form connections with the dopamine-containing dendrites that are developing. Other caveats include whether the phenotype of abnormal striosome-dendron bouquet formation results from the loss of CB1R expression in striosomes, or whether it is an indirect outcome from CB1 loss in another cell type. It is possible that the phenotype is an effect of another perturbation that affects both the afferents and the dendrites. We have not yet been able to determine answers to these crucial issues; further research focused on this neurodevelopmental phenotype could be mapped to a given cell type. Regardless of how CB1R regulates cellular functions and in which cell types, however, the finding that CB1R plays a key role in neurodevelopment of the striatonigral system will need to be taken into account when evaluating phenotypes in mature *Cnr1*^*-/-*^ KO mice and clinically related findings.

### CB1R in striosome-dendron bouquets of adults

Disruption of CB1R function specifically in adults could as well influence striosomal control of nigral dopamine cells, as CB1R is strongly enriched in striosome-dendron bouquets of mature mice (Davis et al., 2018). CB1R signaling in adult mice is reported to control neuronal plasticity in the striatum, midbrain, hippocampus, cerebral cortex, amygdala and cerebellum (Augustin & Lovinger, 2018; Busquets-Garcia, Bains, & Marsicano, 2018). Agonism of presynaptic CB1R can inhibit release of glutamate and GABA from axonal terminals of numerous cell types, including axon terminals from the ventral striatum that disinhibit dopaminergic neurons in the midbrain ventral tegmental area (Riegel & Lupica, 2004; Domenici et al., 2006; Oleson et al., 2014; Tung et al., 2016; Covey et al., 2017; Augustin & Lovinger, 2018). In contrast to striosomal neurons, dopamine-containing neurons express minimal CB1R (Julian et al., 2003). Therefore, CB1R agonism might regulate dopaminergic cell activity via blocking the inhibition and rebound activation from striosomal axons, which are normally mediated by their release of GABA (Lee & Tepper, 2009; McGregor et al., 2019; Evans et al., 2020;). Notable complexities to these proposed effects include that striosomal axons co-express the excitatory slow-acting peptide substance P (Bolam & Smith, 1990; Steiner et al., 1999). Further, CB1R is known to regulate peptide release from non-neuronal cell types (Malenczyk et al., 2013). Whereas activation of CB1R on axon terminals inhibits neurotransmitter release, CB1R activation of astrocytes potentiates glutamate release from surrounding neurons (Navarrete & Araque, 2010). Thus, CB1R can both activate and inhibit neurotransmitter release, depending on its site of action (Stella, 2010). Considering that astrocytes are embedded within bouquet stems (Crittenden et al. 2016), CB1R might regulate bouquet functions by potentiating neurotransmitter release as well as by disinhibition.

Finally, it is notable that striatal expression of CB1R is dysregulated in both Parkinson’s disease (Hurley, Mash & Jenner, 2003; Navarrete et al., 2018; Zeng et al., 1999) and Huntington’s disease (Glass, Dragunow & Faull, 2000; Van Laere et al., 2010), adult-onset disorders with striosomal abnormalities (Crittenden & Graybiel, 2011), dopamine dysregulation and complex motor and mood dysfunction. As a consequence, CB1R has been targeted for preclinical intervention (Kreitzer & Malenka, 2007; Blázquez, et al., 2011; Naydenov et al., 2014). The developmental effects that we report here could yield important clues for work on these and other conditions in which CB1Rs, alone or in conjunction with others, influence transmission in the nigrostriatal dopamine system.

## Acknowledgments

This work was funded by the Broderick Fund for Phytocannabinoid Research at MIT, the Saks Kavanaugh Foundation, NIH/NIMH (R01 MH060379), the Kristin R.Pressman and Jessica J. Pourian ‘13 Fund, Mr. Robert Buxton and William N. & Bernice E. Bumpus Foundation. Special thanks go to Dr. Dan Hu for tissue preparation, to Zhishan Wang for help with histology and cell counting experiments and to Dr. Yasuo Kubota for assistance with manuscript preparation.

---

**Extended Data Figure 2-1.**
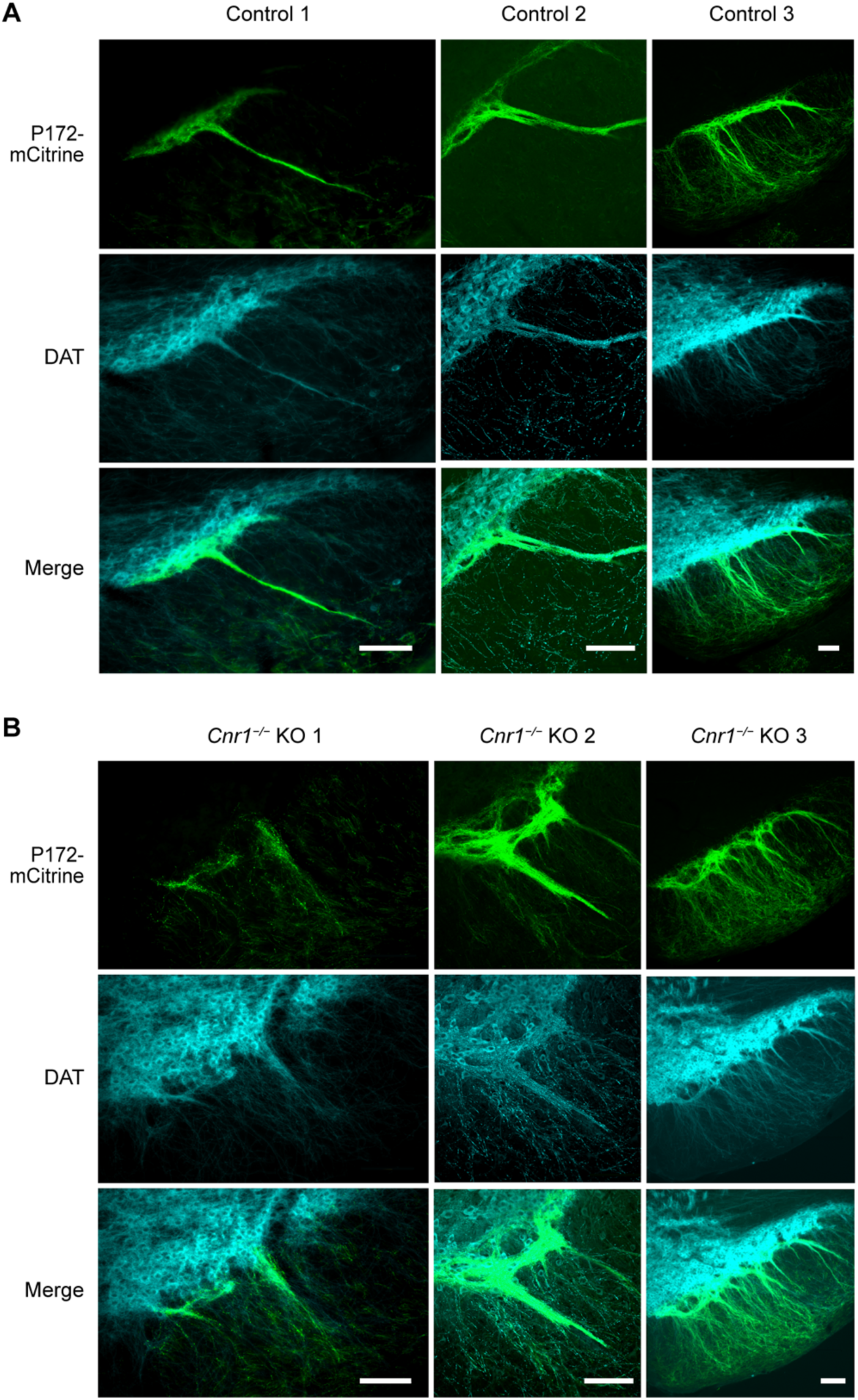
Striosome-dendron bouquet samples in in the SN of controls (***A***) and *Cnr1*^*-/-*^ KO (***B***) mice. Images of the SN from three mice of each genotype showing disorganized and loosely fasciculated striosomal axons (P172-mCitrine fluorescence in green) and dopaminergic dendrites (cyan, DAT immunolabeling in cyan). Coronal sections from the left hemisphere are shown. Scale bars are 100 μm.

**Extended Data Figure 5-1.**
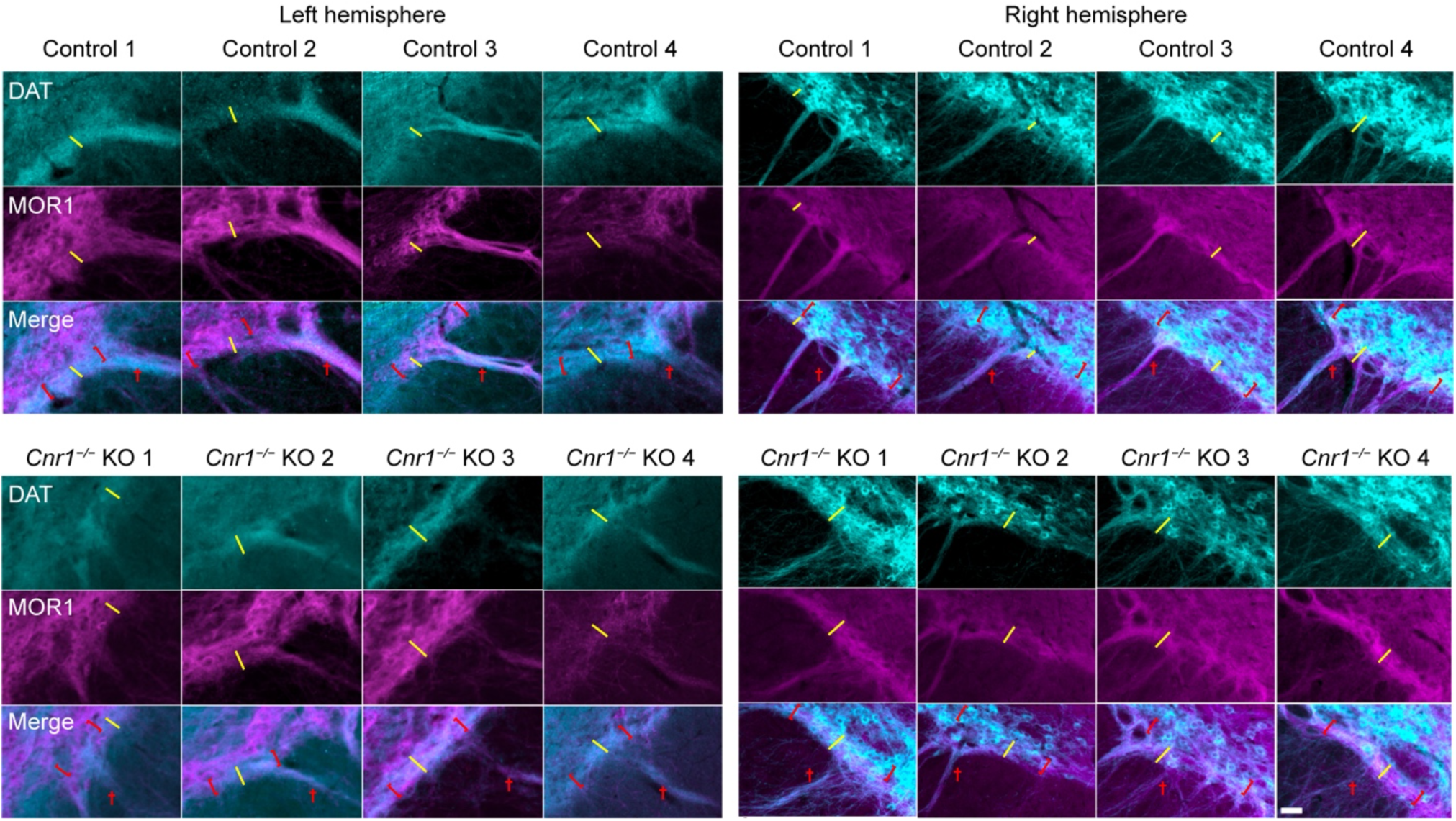
A subset of images from coronal nigral sections used to measure anatomical features in adult *Cnr1*^*-/-*^ KO mice and controls. The SNcv were defined for being immunopositive for both MOR1 and DAT and are delineated by red brackets in the merged image; dendrons are designated by a red cross. Measurements were taken along the white line (one line per section). *N* = 4 mice of each genotype, balanced for sex, *n* = 3 sections per hemisphere for each mouse. Scale bar in the lower right panel is 10 μm and applies to all panels.

**Extended Data Figure 5-2.**
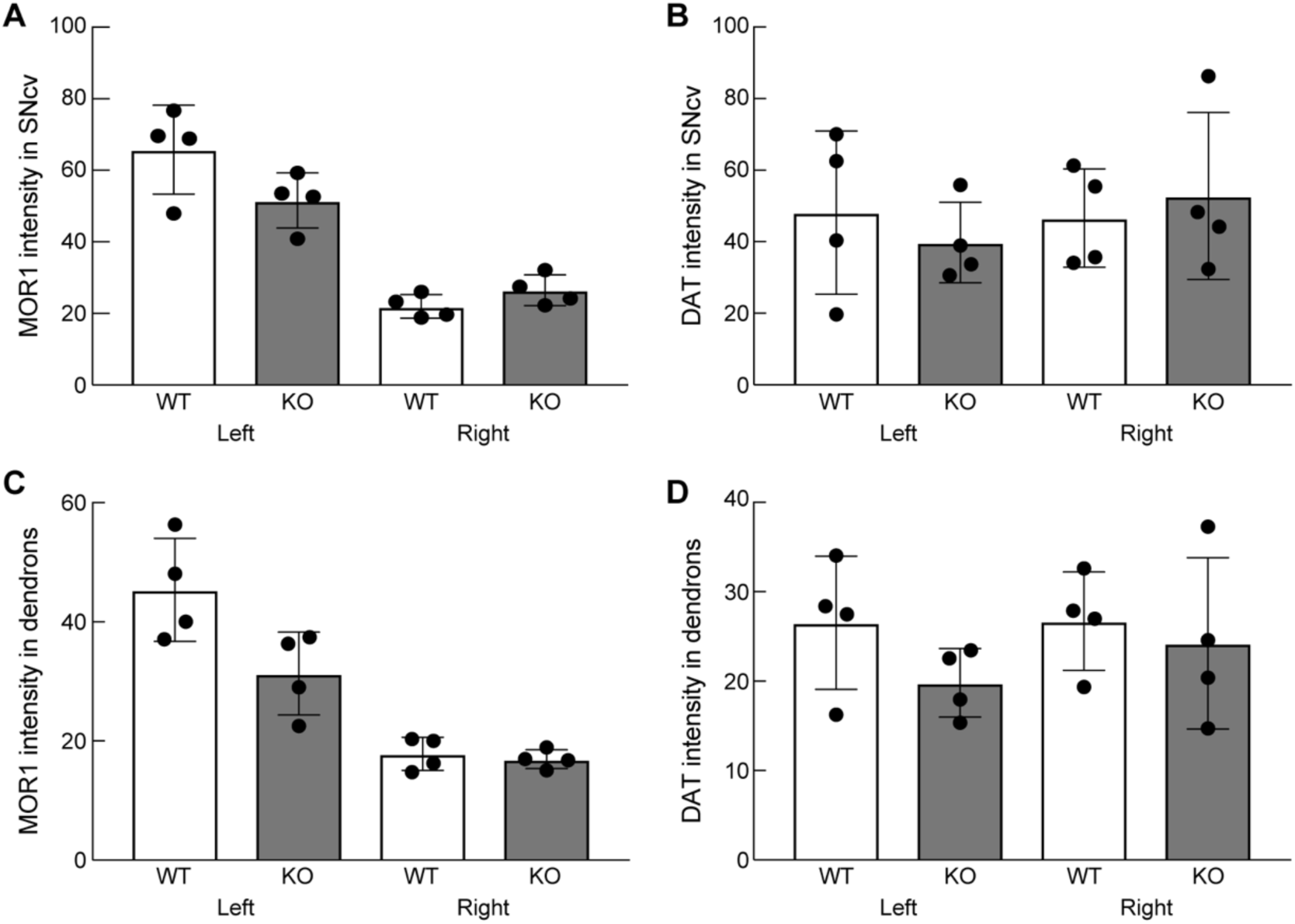
Immunolabeling measurements for MOR1 and DAT in nigral sections from *Cnr1*^*-/-*^ KO mice and controls. There were few significant genotype differences in MOR1 (***A***,***C***) and DAT (***B***,***D***) immunolabeling (arbitrary units) of SNcv (***A***,***B***) or dendrons (***C***,***D***). Genotype comparisons were made only between same-side hemispheres because they were processed together but separately from the opposite hemisphere. (*p* > 0.05 for comparisons between genotypes except for MOR1 immunointensity in left hemisphere dendrons for which *p* = 0.046).

**Extended Data Figure 13-1.**
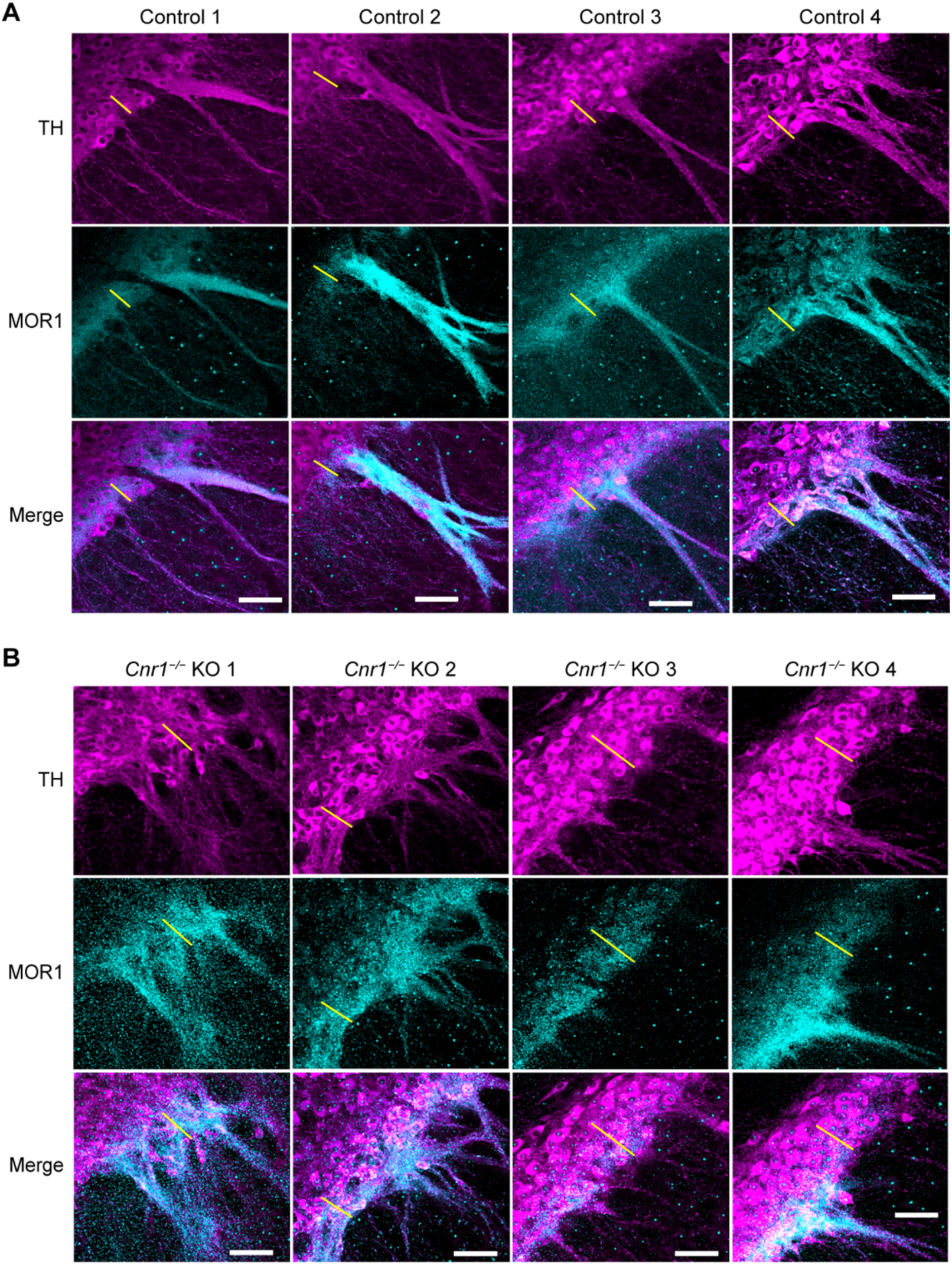
Coronal sections through the left nigra of controls (***A***) and sibling *Cnr1*^*-/-*^ KO (***B***) pups. Co-labeling for TH and MOR1 was used to measure the depth of the SNcv, defined by double-labeled region and designated by yellow line. Scale bars are 50 μm and shown in the merged images panel.

## Notes

### Competing Interest Statement

The authors have declared no competing interest.

